# Maturation stage-specific V-ATPase disassembly can explain neutral pH of lysosome-related organelles

**DOI:** 10.1101/2025.01.13.632790

**Authors:** Ajay Pradhan, Niraj Tadasare, Debolina Sarkar, Vandna Maurya, Lavan K. Bansal, Aaron P. Turkewitz, Santosh Kumar

**Affiliations:** National Centre for Cell Science, NCCS Complex, Savitribai Phule Pune University Campus, Ganeshkhind Road, Pune 411007 Maharashtra, India; Regional Centre for Biotechnology, NCR Biotech Science Cluster, Faridabad 121001 Haryana, India; Department of Molecular Genetics and Cell Biology, The University of Chicago, Chicago IL USA 60637

**Keywords:** LRO, Mucocyst, V-ATPase-a1p, V-ATPase, pHluorin, VPS8ap, Rab7, Secretion, Exocytosis, AP-3

## Abstract

Lysosome-related organelles (LROs) are a heterogeneous family of organelles found in many cell types, whose significant similarities to lysosomes include their acidification by vacuolar-type ATPases (V-ATPases). However, some organelles with hallmarks of LROs are nonetheless non-acidic. Here we investigate this poorly understood phenomenon using the ciliate *Tetrahymena thermophila,* which have prominent secretory LROs called mucocysts. Using three different approaches, we show that mature mucocysts, poised for exocytosis, are non-acidic. Nonetheless, a specific V-ATPase a-subunit is targeted to mucocysts and is indispensable for mucocyst biogenesis and secretion. Consistent with this, immature mucocysts in the cytoplasm appear to be transiently acidified. The stage specificity of acidification can be explained by our finding that several canonical V-ATPase subunits are present in the immature, but not mature, mucocysts. Our data argue that a mucocyst-specific V-ATPase complex is targeted to newly-forming mucocysts and subsequently disassembles at a late stage in *Tetrahymena* LRO maturation, and that this depends on the AP-3 trafficking adaptor.

## Introduction

LROs, which include mammalian Weibel-Palade bodies, melanosomes, acrosomes, basophil granules, Lamellar bodies, and platelet-dense granules, are a family of organelles whose morphological and functional diversity belies that fact that they commonly depend on a set of highly conserved proteins for their biosynthesis (Delevoye et al., 2019; Khawar et al., 2019; Le et al., 2021; Marks et al., 2013; McCormack et al., 2017; Metcalf et al., 2008; Weaver et al., 2002). Though sharing features with both late endosomes and lysosomes, LROs are functionally, morphologically, and/or compositionally distinct (Luzio et al., 2014; Huizing et al., 2008; Marks et al., 2013). They are formed by a multistep process in which an immature organelle matures by acquiring cargo from both secretory pathways as well as traffic from endosomes (Bonifacino, 2004; EC et al., 2000; Raposo et al., 2007). Many of the protein cargoes targeted to LROs within the endomembrane network rely on receptors and cytoplasmic adaptors for their selection as well as complexes to tether and fuse vesicles (Banushi and Simpson, 2022; Marks et al., 2013). These include the AP-3 adaptor, the homotypic fusion and protein sorting (HOPS) complex, the small GTPases Rab32 and Rab38, the sorting receptor sortilin/VPS10, and three biogenesis of LROs (BLOC) complexes (Banushi and Simpson, 2022; Raposo et al., 2007; Marks et al., 2013; Gautam et al., 2006; Hermey, 2009; Starcevic et al., 2002; Wasmeier et al., 2006; Kaur et al., 2017). Phylogenetic analysis of LRO machinery revealed that these LRO components were present in the last eukaryotic common ancestor (LECA) and a basic element of eukaryotic membrane trafficking (More et al., 2024).

One cargo broadly shared among LROs is the Vacuolar-type proton ATPase (V-ATPase) complex, whose presence drives acidification of the LRO lumen. Acidification plays a critical role for many types of LROs (EC et al., 2000; Pejler et al., 2017; Matsumoto and Nakanishi-Matsui, 2019; Sun-Wada et al., 2003). For instance, T lymphocytes and natural killer cells synthesise LROs called lytic granules, that maintain an acidic pH essential for the targeted release of macromolecules to eliminate virally-infected or tumor cells (Griffiths and Argon, 1995). Type II alveolar epithelial cells synthesise lamellar bodies that store lung surfactant. The processing and therefore release of surfactant proteins from lamellar bodies requires an acidic pH (Weaver et al., 2002; Chander et al., 1986; Chintagari et al., 2010). Sperm cells synthesize the acrosome, which contains various hydrolytic enzymes that are secreted to help the sperm in penetrating the egg’s protective layers. An acidic environment is essential for the biogenesis and exocytosis of the acrosome (Berruti and Paiardi, 2011). In endothelial cells, Weibel-Palade bodies are formed from the trans-Golgi network (TGN) and mature into elongated, cigar-like structures that store the adhesive glycoprotein von Willebrand factor (VWF). This glycoprotein is essential for facilitating interactions between platelets and the vascular wall. The luminal acidification of Weibel-Palade bodies is a key factor for the dense packing of VWF and its release from endothelial cells (Naß et al., 2021).

V-ATPases are well-studied complexes that are composed of two macromolecular domains, V_1_ and V_0_. V_1_, the catalytic domain, is a 650-kDa peripheral complex comprised of A_3_B_3_CDE_3_FG_3_H subunits, that hydrolyzes ATP to generate the energy required for pumping protons (Forgac, 2007; Futai et al., 2019; Breton and Brown, 2013). V_0_ is a 260-kDa integral membrane protein complex that forms a proton channel; it consists of ac_8_c′c′′def subunits (Breton and Brown, 2013; Forgac, 2007; Futai et al., 2019). Thus, both the V_1_ and V_0_ subunits are required to form a fully functional holo-V-ATPase complex. Since V-ATPases are active in diverse intracellular compartments, compartment-specific stimulation of assembly or disassembly may be one organizing principle of the endomembrane network. For example, increasing assembly has been proposed as a cause of increased acidity throughout the endocytic route in mammalian cells (Lafourcade et al., 2008). Positive regulators of V-ATPase complex assembly include the heterotrimeric chaperone complex “RAVE (Regulator of ATPases of Vacuolar and Endosomal membranes)” as well as glycolytic enzymes phosphofructokinase-I and aldolase (Parra and Kane, 1998; Xu and Forgac, 2001; Lu et al., 2007; Chan and Parra, 2014; Smardon et al., 2015). Conversely, the complex’s enzymatic activity can be downregulated by disassembly, in which the V_1_ domain separates from the membrane-integral V_0_ domain (Parra et al., 2014; Oot et al., 2017). In yeast and mammalian cells, reversible V-ATPase assembly and disassembly have been reported in response to stimuli including glucose and amino acid starvation and refeeding, and oxidative stress (Chan and Parra, 2014; Trombetta et al., 2003; Sautin et al., 2005; Stransky and Forgac, 2015; Bodzęta et al., 2017; Parra and Kane, 1998; Khan et al., 2022).

In addition to compartment-specific assembly/disassembly, V-ATPase activities in endocytic and secretory pathways can be tailored for different compartments based on the expression of multiple paralogs for some subunits (Toei et al., 2010). These paralogs have historically been called isoforms in the field, a terminology we will therefore adopt in this Introduction. Such isoforms can, among other things, provide determinants for compartment-specific targeting. In yeast, subunit-a has two isoforms, encoded by the *VPH1* and *STV1* genes (Manolson et al., 1994, 1992). V-ATPases containing these alternate a-subunits differ in proton pump energy-coupling efficiency, subcellular localization, and in the regulation of subunit association under cellular metabolic conditions (Kawasaki-Nishi et al., 2001; Manolson et al., 1994). In mammals, most V-ATPase subunits exist in multiple isoforms that are expressed in a tissue-specific manner, and isoform-specific activities or localization have been demonstrated in many species (Toei et al., 2010; Breton and Brown, 2013; Forgac, 2007; Chu et al., 2021). For example, there are four subunit-a isoforms (a1-a4) which direct their cognate complexes to distinct cellular destinations (Sun-Wada and Wada, 2013; Chen et al., 2024; Matsumoto et al., 2018; Toei et al., 2010; Sun-Wada and Wada, 2022; Sun-Wada et al., 2007). Isoforms a1, a2, and a3 are associated with coated vesicles, the Golgi apparatus/early endosomes, and late endosomes/lysosomes, respectively (Breton and Brown, 2013; Chu et al., 2021; Toei et al., 2010). Subunit a3 in pancreatic islet cells also localizes to insulin granules, which are LROs whose maturation requires luminal acidification (Sun-Wada et al., 2006). An extreme example of such isoform-specific targeting comes from the ciliate *Paramecium tetraurelia,* in which seventeen a-subunit isoforms localize to at least seven different compartments (Wassmer et al., 2006). Compartment-specific isoforms are also known to play additional roles. For example, mammalian a3 is a determinant for Rab7-dependent trafficking of LRO proteins in osteoclasts (Matsumoto et al., 2018, 2022; Nakanishi-Matsui et al., 2024). Notwithstanding the importance of compartment-specific optimization, the conserved activity of V-ATPases is proton pumping. However, several recent studies have identified non-acidic and nondegradative LROs across different cell types, including melanocytes and neurons, as well as in unicellular eukaryotes such as *Dictyostelium* (Padh et al., 1993; Fok et al., 1993; Cheng et al., 2018; Hurbain et al., 2018; Correia et al., 2018; Yap et al., 2018). For example, LROs in melanocytes called melanosomes are only transiently acidic: premelanosomes are acidic, whereas mature melanosomes are non-acidic, which may be more consistent with their role as storage rather than degradative compartments (Hurbain et al., 2018; Correia et al., 2018; Le et al., 2021). A similar cellular logic may underlie the LAMP1-positive but non-acidic and non-degradative (i.e., lacking acid hydrolases) endolysosome-related organelles (ELROs) organelles found in neuronal cells (Cheng et al., 2018; Yap et al., 2018). In *Dictyostelium discoideum*, LROs called post-lysosomes originate as acidic lysosomes but subsequently lose their acidity and mature into neutral pH organelles (Padh et al., 1993; Fok et al., 1993). The mechanisms involved in the neutralization of these organelles in widespread eukaryotic lineages are not known.

One organism with non-acidic LROs is the ciliate *Paramecium tetraurelia*, where secretory LROs called trichocysts dock at the plasma membrane to release their contents via regulated exocytosis in response to extracellular stimuli (Plattner, 2017; Lumpert et al., 1992; de Loubresse et al., 1994). However, V-ATPase activity nonetheless appears required for trichocyst formation. To understand this paradox and shed light on the nature of neutral pH LROs, we have extended the analysis in *Paramecium* using the related Oligohymenophorean ciliate *Tetrahymena thermophila*, where the homologous organelles to trichocysts are known as mucocysts (Turkewitz, 2004). These have been characterized as LROs based on multiple lines of molecular evidence. Mucocysts, like LROs in many organisms, undergo biochemical and morphological maturation (Briguglio et al., 2013; Kumar et al., 2014, 2015; Kaur et al., 2017; Sparvoli et al., 2018). The most abundant mucocyst cargo proteins are encoded by two multigene families: *GRL* (granule lattice) and *GRT/IGR* (granule tip/induced on granule regeneration) (Bowman et al., 2005; Cowan et al., 2005; BOWMAN et al., 2005). Grls are synthesized as proproteins and subsequently cleaved and trimmed by processing enzymes Cth3p (Cathepsin 3) and Cth4p to generate mature Grls (Kumar et al., 2015, 2014). Mature Grls assemble to form a crystal that fills the mucocyst lumen and expands upon exocytic fusion of a docked mucocyst, propelling the contents out to the extracellular space (Cowan et al., 2005). Grt/Igr proteins are not required for crystal formation or for content propulsion, and at least some family members are sub-localized within the mucocyst lumen (BOWMAN et al., 2005).

Importantly, cargo delivery to mucocysts involves at least two different pathways. The delivery of Cth3p and Grt/Igrs, but not Grls, requires the sortilin/VPS10 receptor Sor4p and an endosomal SNARE, Stx7l1p (Syntaxin 7-like) (Briguglio et al., 2013; Kaur et al., 2017; Sparvoli et al., 2018). The Grl and Grt proteins localize to different vesicles during mucocyst formation; these and other data argue that heterotypic fusion between secretory and endosomal vesicles is a key step in formation of ciliate LROs, like in other lineages. In *Tetrahymena* that fusion requires VPS8a, a subunit of a dedicated CORVET (class C core endosomal vacuole tethering) complex (Sparvoli et al., 2018). The formation of mucocysts also requires the AP3 complex, and cells lacking the AP3µ subunit accumulate abortive mucocyst intermediates distinct from those in *VPS8A* knockout cells (Kaur et al., 2017). However, the presumed AP3-dependent mucocyst cargo is not known.

In this study, we confirmed that mature mucocysts are not acidified. Nonetheless, their biogenesis and more specifically maturation depends on the V-ATPase a1-subunit isoform, one of six a-subunit paralogs/isoforms in this species, and which is localized to both immature cytoplasmic mucocysts as well as mature docked mucocysts. By targeting a pH-sensor to the mucocyst lumen, we found evidence the immature mucocysts are transiently acidified. Consistent with this, we found that other subunits of the V-ATPase are also present at that stage in mucocyst formation. Analysis of cells lacking a1-subunit expression suggests that the failure to acidify during mucocyst maturation leads to predictable defects in proprotein (Grl) processing, but also defects in the heterotypic vesicle fusion characteristic of LRO formation. In striking contrast, several other V-ATPase subunits are absent from mature docked mucocysts. Our findings suggest that the V-ATPase complex disassembles during mucocyst maturation, and can therefore account for the neutral pH of the mature organelles. In cells lacking AP3µ, the accumulated mucocyst intermediates appear to retain the full cohort of V-ATPase subunits, indicating that the activity of AP3 is required for V-ATPase disassembly.

## Results

### Expression profiling reveals co-regulation of mucocyst-associated proteins and V-ATPase-a1 subunit of the V_0_ domain of V-ATPase complex in *T. thermophila*

The *T. thermophila* macronuclear genome has more than 24,000 predicted genes, including 11 that encode the subunits of the V_1_ domain and 12 that encode the subunits of the V_0_ domain of the V-ATPase complex (Coyne et al., 2008; Eisen et al., 2006). To ask whether there might be paralog-encoded subunits specialized for mucocysts, we followed up on earlier findings that many genes encoding mucocyst-related proteins are highly coregulated under a variety of conditions (Xiong et al., 2013; Briguglio et al., 2013; Kumar et al., 2014; Miao et al., 2009; Xiong et al., 2011). We used the online tools at the Tetrahymena Functional Genomics Database (http://tfgd.ihb.ac.cn/) to identify V-ATPase subunits that are coregulated with known mucocyst-associated genes. This search revealed that TTHERM_01332070, encoding an a-subunit of V-ATPase V_0_, was coregulated with known mucocyst-associated genes **(Figure 1A)** and therefore represented a strong candidate for a mucocyst-specific V-ATPase subunit. We have given this paralog the name a1. In addition to TTHERM_01332070, the *T. thermophila* genome contains five additional paralogs, which we named *a2* (TTHERM_00463420), *a3* (TTHERM_00266570), *a4* (TTHERM_00372530), *a5* (TTHERM_01005210) and *a6* (TTHERM_01014710). These paralogs are significantly diverged, with a maximum of 45% amino acid sequence identity to the a1 paralog **(Figure S1)**, and they show transcriptional profiles distinct from that of a1 **(Figure 1B)**, consistent with the idea that they belong to complexes specialized for other cellular compartments.

**Figure 1.**
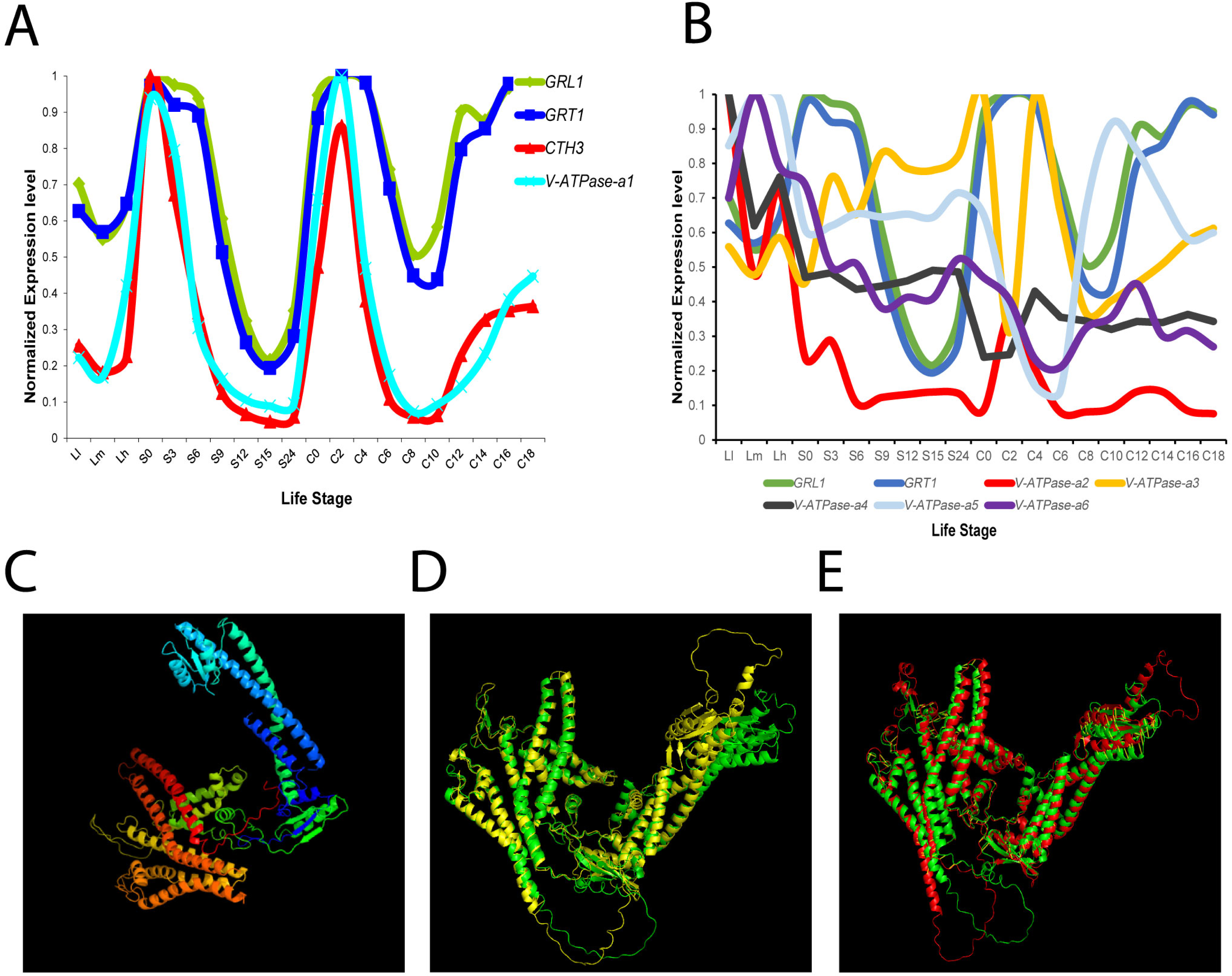
(A) Expression profiling identifies a subunit of the V*-*ATPase complex (V-ATPase-a1) that is highly coregulated with known mucocyst associated genes (*GRL1*, *GRT1* and *CTH3*). Transcript data were retrieved from the Tetrahymena Functional Genomics Database (TFGD). For plotting the profiles, each value was normalized to the respective gene’s maximum expression level. (B) Other paralogs of the *V-ATPase-a* subunit (*a2-a6*) are not coregulated with known mucocyst genes. (C) The predicted structure of V-ATPase-a1p of *Tetrahymena* by the web portal Phyre2. The image colors are ordered like the rainbow, from the N to the C terminus. (D-E) The predicted structure of V-ATPase-a1p of *Tetrahymena thermophila* (green), *Saccharomyces cerevisiae* (yellow) and *Mus musculus* (red) by AlphaFold structure prediction. V-ATPase-a1p structure of *Tetrahymena thermophila* superimposed with V-ATPase-a1p of *Saccharomyces cerevisiae* (D) and *Mus musculus* (E) by PyMOL software.

*Tetrahymena* V-ATPase-a1p (where “p” denotes the protein) shares ∼30% sequence identity with human subunit-a isoforms (a1p: ATP6V0A1, a2p: ATP6V0A2, a3p: ATP6V0A3 and a4p: ATP6V0A4). The predicted structure of the *T. thermophila* V-ATPase-a1p was strikingly similar to those of V-ATPase-a subunits in other lineages (shown for Opisthokonts) with 100% confidence and 85% coverage **(Figure 1C).** Structures similar to the *T. thermophila* V-ATPase-a1p (green) in the AlphaFold Protein Structure Database included the homologous a-subunits from *Saccharomyces cerevisiae* (yellow; A0A7I9G792) and *Mus musculus* (red; Q9Z1G4), the former with an 84% per-residue model confidence score (pLDDT) **(Figure 1D-E)**. The phylogenetic mapping of the a-subunits in *Tetrahymena* and in other organisms is consistent with the results of the structural analysis **(Figure S2)**.

### V-ATPase-a1p localizes to mucocysts

To ask whether the *Tetrahymena* a1 subunit, implied via coregulation to be associated with mucocysts, is physically associated with this compartment, we tagged the protein via C-terminal fusion of the endogenous gene with 2 x mNeon. The expression in *Tetrahymena* of a fusion protein of the expected size was confirmed by immunoprecipitation followed by Western blotting **(Figure 2A)**. The localization of the fluorescent protein was examined by confocal microscopy and compared with mucocyst markers. In live cells, V-ATPase-a1p localized to linear arrays of fluorescent puncta at the cell surface, consistent with its accumulation in mucocysts **(Figure 2B, left panel, inset and Movie S1)**. In cross sections of the same cells, docked mucocysts are seen as elongated vesicles **(Figure 2B, right panel, inset and Movie S1)**. In addition, a significant fraction of V-ATPase-a1p localizes to heterogeneous cytoplasmic puncta at some distance from the cortex (which we call “non-cortical”). Approximately 75% of V-ATPase-a1p is present in the non-cortical compartments, with the remaining quarter located at the cell cortex in the docked mucocysts (**Figure 2C-D**).

**Figure 2.**
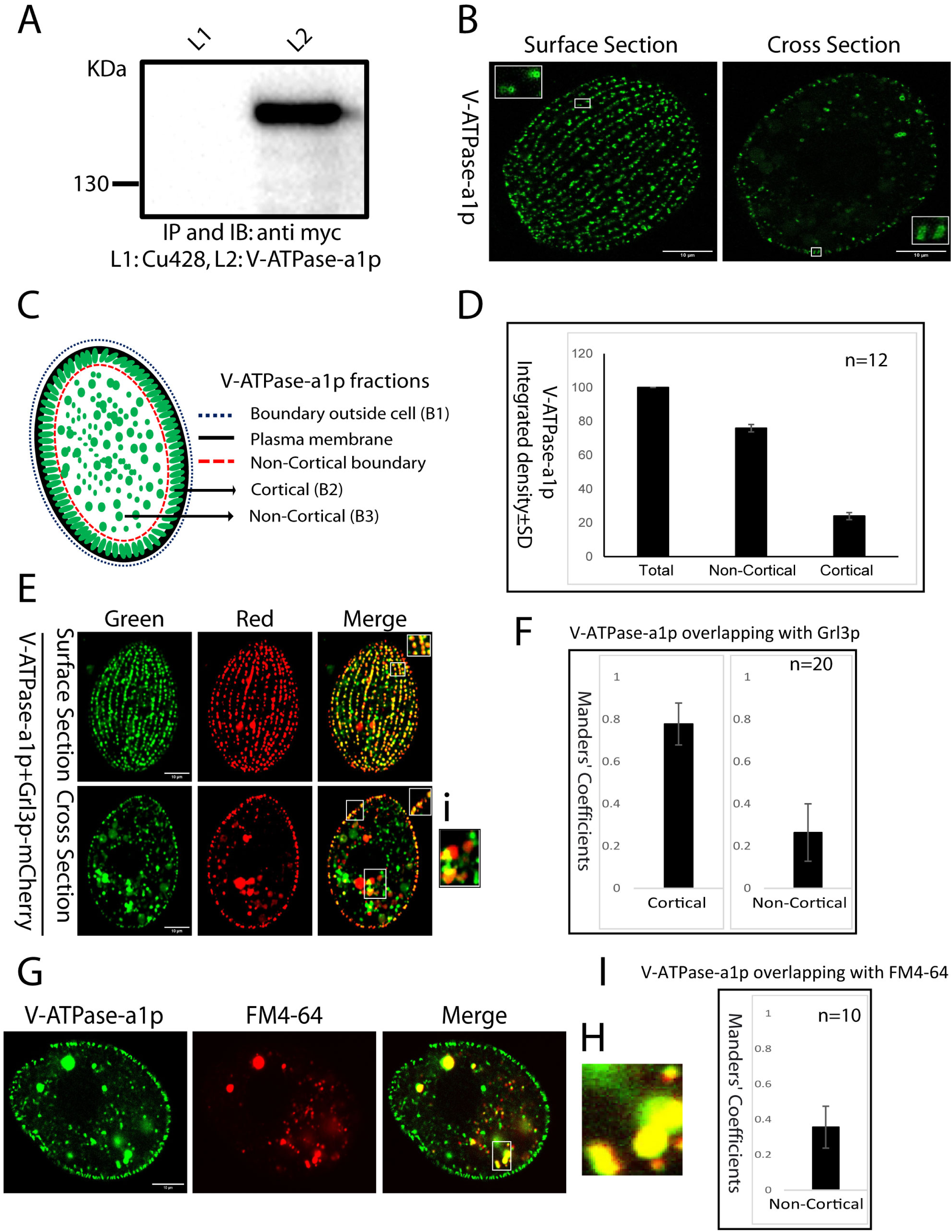
Expression and localization of V-ATPase-a1p. (A) Detergent solutes of cells expressing V-ATPase-a1p were immunoprecipitated using anti-c-Myc Agarose beads. Pull down samples were separated by SDS-PAGE and blotted with mouse monoclonal anti-c-myc antibody. Blots were incubated with anti-mouse-IgG-HRP for IP and developed by SuperSignal West Femto Maximum Sensitivity Substrate. The expected size-specific band was only found in the V-ATPase-a1p lane. (B) In optical surface and cross section (left and right, respectively), images of cells endogenously expressing 2xmNeon-6c-myc-tagged V-ATPase-a1p subunit were acquired by super resolution confocal microscopy with low exposure conditions. At the cell surface, linear arrays of fluorescent puncta correspond to docked mucocysts, which show up in cross sections of the same cells as elongated vesicles. The inset reveals that V-ATPase-a1p accumulates in these docked mature mucocysts. (C) Cartoon representing localization of V-ATPase-a1p at cortical and non-cortical compartments of cells. (D) The mean fluorescence intensity associated with cell (B1), cortical (B2), and non-cortical (B3) was measured and converted to integrated density. Most V-ATPase-a1p is found in non-cortical compartments. (E) Cells co-expressing V-ATPase-a1p-2xmNeon with mucocyst marker Grl3p-3xmCherry, both at endogenous loci. V-ATPase-a1p and Grl3p overlap extensively at docked mucocysts. Images are shown as single slices, for clarity. The inset Ei depicts co-localization of V-ATPase-a1p with Grl3p. To measure co-localization, the images were processed to remove noise and adjust the thresholds, with co-localization quantified using the Fiji-BIOP JACoP plugin. (F) 20 non-overlapping images from panels E were used to calculate the percentage of overlap (Mander’s coefficient) between V-ATPase-a1p and Grl3p in cortical (surface section) and non-cortical (cross section) domains. The error bars show the SDs. (G) V-ATPase-a1p in non-cortical puncta shows some overlap with endosomal marker FM4-64. Cells expressing V-ATPase-a1p-2xmNeon were incubated for 5 min with 5μM FM4-64 and then washed with 10mM Tris media (pH 7.4). Images were acquired after 30-40 minutes. (H) The inset (H) is from panel 2G. (I) 10 non-overlapping images from panels 2G were used to calculate the percentage of overlap (Mander’s coefficient) between V-ATPase-a1p-2xmNeon and FM4-64 in non-cortical puncta. Scale bars, 10 μm.

Localization of a1p to mucocysts was confirmed by expressing tagged a1p in cells co-expressing mCherry-tagged mucocyst protein Grl3p. The two proteins showed strong co-localization (Manders’ Coefficients 0.78± SD 0.10) in the docked mature mucocysts at the cell cortex **(Figure 2E, inset and 2F, left)**. In optical cross sections of the same cells, more limited overlap (Manders’ Coefficients 0.26± SD 0.14) between a1p and Grl3p was observed, corresponding to overlap in non-cortical structures (**Figure 2E, Ei inset and 2F, right)**. We interpret non-cortical structures containing Grl3p as likely vesicular intermediates in mucocyst biogenesis.

Our results also indicate that a large fraction of non-cortical a1p is associated with compartments that do not also contain Grl3p. We found that non-cortical a1p showed significant overlap with the endocytic tracer FM4-64 (Manders’ Coefficients 0.36± SD 0.12) (**Figure 2G-I**) and lesser overlap with the late endosomal marker Rab7p (Manders’ Coefficients 0.17± SD 0.04) **(Figure S3A-C)** or with Lysotracker (Manders’ Coefficients 0.08± SD 0.04) **(Figure S3D-F)**. Collectively, the findings support the idea that non-cortical V-ATPase-a1p is associated with biosynthetic mucocyst intermediates, and with endosomes that may be primarily be considered early or recycling.

Since other V-ATPase-a1 paralogs in *T. thermophila* show very different transcriptional profiles, we predicted that they would also reside in different compartments from a1. To examine this, we analyzed the localization of V-ATPase-a2p. This protein was found to be localized mostly near the plasma membrane but not co-localized with the mucocyst marker Grl3p **(Figure S3G)**.

### Mucocysts are not stably acidic in *Tetrahymena*

*Paramecium* trichocysts which are homologous to *Tetrahymena* mucocysts, are not detectibly acidic (de Loubresse et al., 1994; Lumpert et al., 1992). However, in *Tetrahymena* mucocysts the key protease responsible for proGrl cleavage, called Cth3p, is related to lysosomal enzymes (Kumar et al., 2014), and expected to be most active under acidic conditions. Therefore, we asked whether mucocysts maintained acidic luminal conditions. First, we incubated cells expressing Igr1p-mCherry with Acridine orange (AO) dye, which accumulates in acidic vesicles. There was no visible accumulation of AO at docked mature mucocysts **(Figure 3A)**. Similarly, we incubated cells expressing Grl3p-mCherry with an alternative probe for acidic compartments, Protonex Green (Prakash et al., 2021). Consistent with the AO results, we observed no concentration of Protonex Green in mature mucocysts at the cell cortex **(Figure 3B)**.

**Figure 3.**
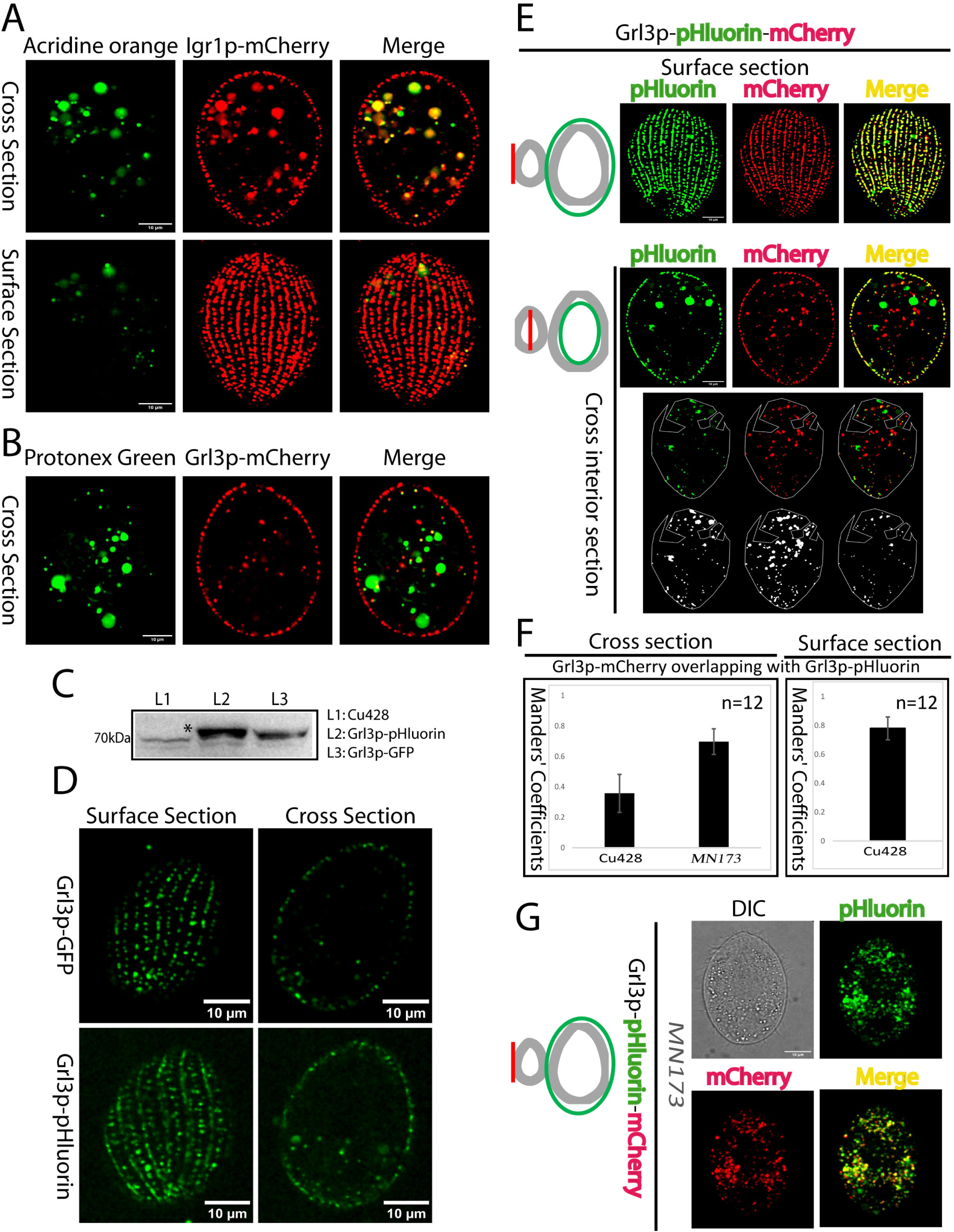
The mucocyst lumen is not stably acidic. (A) Cells expressing mucocyst cargo Igr1p-3xmCherry incubated with Acridine Orange (AO). There is minimal overlap between AO and Igr1p in docked mucocysts. Confocal images are shown as single slices, corresponding to cross or surface sections, for clarity. (B) Protonex Green incubated with cells expressing mucocyst cargo Grl3p-3xmCherry. No significant co-localization is seen between Protonex Green and Grl3p in docked mucocysts. (C) Cell lysates (60,000 cell equivalents) were separated using SDS-PAGE, transferred to PVDF and probed with anti-GFP mAb, which also recognizes pHluorin. Bands corresponding to Grl3p-GFP and Grl3p-pHluorin are indicated by an asterisk. (D) Grl3p-GFP and Grl3p-pHluorin constructs were expressed at the native *GRL3* locus. The Super ecliptic pHluorin (SEP) only emits fluorescence at pH 6.5 or higher. Both Grl3p-GFP and Grl3p-pHluorin show strong signals in docked mucocysts, consistent with their characterization as a non-acidic compartment. (E-G) Endogenous expression of Grl3p-pHluorin-mCherry in Cu428 (wild type) and mutant MN173 cells. (E) In wild type cells, dual emission from both pHluorin and mCherry at each fluorescent spot was measured as co-localization, and was analyzed in cell surface (top) and cross-sections (middle). The images were processed to remove noise and adjust thresholds, with co-localization quantified using the Fiji-BIOP JACoP plugin. The edges of the cross-sections were excluded (marked by a green boundary line in the illustration on the left) to restrict analysis to non-cortical structures. To minimize false co-localization due to the bright autofluorescence from large food vacuoles, we selected areas lacking food vacuoles (bottom) for quantitative analysis. The color images at the bottom illustrate co-localization in selected non-cortical regions, while the grey images reveal co-localization after threshold adjustments for the green and red channels. Surface sections were analyzed to measure co-localization in cortical structures. The pHluorin and mCherry signals showed complete overlap within the docked mucocysts. There was also significant co-localization of red and green signals in non-cortical structures, as shown in cross-sections of the same cells. (F) 12 non-overlapping images from panels E and G were analysed for both non-cortical (left) and cortical puncta (right). (G) Co-localization was measured in cross-sections of MN173 cells, as shown in panel E. Significant overlap of green and red signals was observed in the non-cortical region. The error bars show the SDs. Scale bars, 10 μm.

Given the significance of these data for interpreting the role of V-ATPase-a1, we took a third and more flexible approach to analyzing the pH of mucocysts by endogenously tagging Grl3p with either mEGFP or the super ecliptic pHluorin (SEP) variant, a green fluorescent protein whose emission is relatively quenched at acidic pH. In mammalian cells in which SEP is targeted to the lumen of acidified secretory vesicles, SEP emits bright fluorescence only after the vesicles undergo exocytosis, when their contents become neutralized (Benčina, 2013; Martinière et al., 2013; Michaluk and Rusakov, 2022). In our experiments, the salient advantage to using tagged Grl3p as a pH sensor is that it also allowed us to analyze the pH in the lumen of non-docked, immature mucocysts.

We first confirmed the expression of the fusion proteins Grl3p-mEGFP and Grl3p-pHluorin by western blotting using anti-GFP mAb **(Figure 3C)**. Importantly, by confocal microscopy we detected strong fluorescence signals for both Grl3p-mEGFP and Grl3p-pHluorin in docked mucocysts **(Figure 3D)**. Since the pHluorin signal in the easily identified docked mucocysts was not quenched, this result provides additional confirmation that mature mucocysts are not acidified. Since we could also detect non-quenched cytoplasmic pHluorin signal in non-cortical puncta, immature mucocysts may also be non-acidic.

To address the pH of mucocyst intermediates more rigorously, we endogenously tagged Grl3p in tandem with pHluorin followed by mCherry, whose fluorescence emission is not pH sensitive (Shaner et al., 2004, 2005; Doherty et al., 2010; Hundeshagen et al., 2011). By simultaneously collecting data in the red and green channels, we could detect acidic mucocysts (or mucocyst intermediates) as puncta showing red but not green fluorescence. In the docked mucocysts of wildtype cells (i.e., the cortical field), the large majority of red puncta also showed green fluorescence, with an overlap between signals of (Manders’ Coefficients 0.78± SD 0.088) **(Figure 3E, top panel and 3F, right panel)**. In contrast, the overlap between green and red signals was significantly lower in non-cortical puncta (Manders’ Coefficients 0.36± SD 0.13) **(Figure 3E, middle and bottom panels, and 3F, left panel)**. Thus, there is a significant population of Grl3p-containing, mucocyst-related puncta in the cytoplasm in which the pHluorin emission is quenched. This result strongly suggests that mucocysts are acidified at some stage in their formation.

A collection of mutant cell lines in *Tetrahymena* have characterized defects in mucocyst biogenesis or exocytosis. One such lines is MN173, in which the mucocysts undergo normal biochemical and morphological maturation but fail to dock at the plasma membrane and thus remain primarily dispersed in the cytoplasm (Chilcoat et al., 1996; Melia et al., 1998). We therefore expected that Grl3p-pHluorin-mCherry, expressed in MN173 cells, would produce non-cortical puncta with a strong overlap of green and red signals, from the non-acidic mature cytoplasmic mucocysts. Indeed, such cells showed complete co-localization of pHluorin and mCherry in the cytoplasm, supporting the validity of our approach **(Figure 3F, left panel and 3G)**. Taken together, our results strongly suggest that mucocysts are transiently acidified during their formation, while mature mucocysts are non-acidic.

### The holo-V-ATPase complex does not persist during mucocyst maturation

Our localization data for V-ATPase a1, shown above, are consistent with the idea that this subunit is present in the transiently acidified immature non-cortical mucocysts, but also clearly demonstrate that the subunit is still present in non-acidified mature docked mucocysts. We therefore asked whether other subunits of the V-ATPase complex were similarly localized, i.e., whether the V-ATPase is stably maintained during mucocyst maturation, or whether partial disassembly occurs so that some subunits are missing in mature mucocysts. To avoid the potential ambiguities that arise from studying genes that have paralogous copies, we focused on V-ATPase subunits encoded by single genes in *T. thermophila*. To begin, we used approaches including BLAST and structural homology to reveal that putative V_1_ domain subunits in *Tetrahymena*, including Vma1p (subunit-A), Vma8p (subunit-D) and Vma10p (subunit-G) are expressed as single genes **(Table S1 and Figure S4A-E)**. The identity of these genes was determined using multiple approaches. The *T. thermophila* homologs for Vma1p, Vma8p and Vma10p) were found to have high sequence identity via BLAST with Vma subunits in the Opisthokont *S. cerevisiae* **(Table S1, Figure S4A-E)**. Consistent with the sequence identity, the Phyre2-predicted structures of Vma8p and Vma1p revealed high structural similarity with subunit-D and subunit-A of the V_1_ domain of mammalian cells, respectively **(Figure S4A and S4B)**. *T. thermophila* Vma10p also showed predicted structural similarity to a V-ATPase subunit in *S. cerevisiae*, subunit-G, though this was more limited (62% coverage and 28% structural identity) **(Figure S4C)**. The predicted structural similarities between *T. thermophila* and human V-ATPase subunits were also supported by AlphaFold predictions, for subunit-D (**Figure S4D)** and subunit-A (**Figure S4E)** of *T. thermophila* (green) and human (blue). Taken together, these sequence and structural similarities strongly argue that these genes in *T. thermophila* encode components of the proton-translocating V-ATPase complex.

For each of these three V-ATPase subunits in *Tetrahymena*, each encoded by a single gene without paralogs, we then asked whether they were present in mucocysts. We imaged cells that endogenously co-expressed mCherry-tagged mucocyst marker Grl3p with mNeon-tagged Vma1p, Vma8p or Vma10p **(Figure 4A-J)**, and measured co-localization using the BIOP JACoP plugin in surface **(Figure 4A)** and cross sections **(Figure 4B)**. We found negligible overlap between the Vma subunits and Grl3p at the cell cortex in mature mucocysts **(Figure 4C, 4E, 4G and 4I)**. As expected, based on results shown above, there was instead complete overlap between Grl3p and V-ATPase-a1 at the cell cortex **(Figure 4K)**.

**Figure 4.**
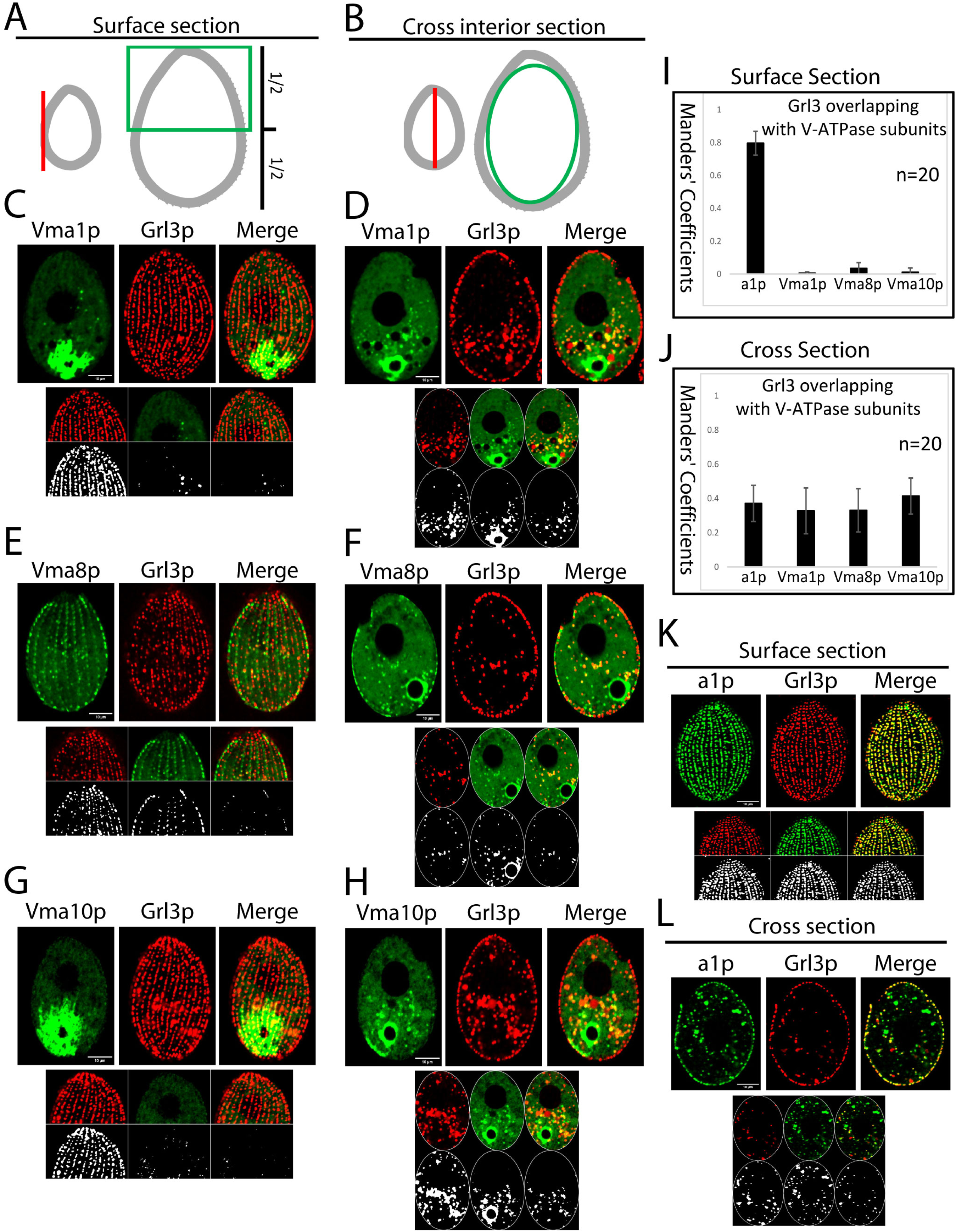
The V-ATPase complex in mature, but not immature, mucocysts lacks multiple subunits. (A-B) The co-localization of Vma subunits and Grl3p was measured using cell surface (A) and cross (B) sections. For the former, the edge of the cross-sections was excluded (indicated by green boundary line in cartoon, panel B) in order to measure colocalization in non-cortical structures. The surface sections (panel A) were used to measure colocalization in cortical structures. To reduce false co-localization in cortical sections due to the bright signal from the contractile vacuole near the cell posterior, only the anterior ∼half of the cell was analyzed. (C-H) Cells co-expressing 2xmNeon-tagged Vma1p (C-D), Vma8p (E-F), and Vma10p (G-H) with the mucocyst marker Grl3p-3xmCherry, all at their endogenous loci. In both cross and surface sections, Vma subunits are localized at the contractile vacuoles (CVs) and in non-cortical puncta, as well as at the cell cortex. Co-localization was measured using the Fiji-BIOP JACoP plugin, as shown in panel A and B. The color images at the bottom show co-localization in selected non-cortical (cross section) and cortical compartments (surface section). Grey images show co-localization after adjusting the thresholds for the green and red channels. There was no co-localization of Vma subunits with Grl3p in mature, docked mucocysts at the cell cortex. In contrast, there was significant overlap between Grl3p and the Vma subunits in non-cortical compartments. (I-J). 20 non-overlapping images from panels C-H, K and L were used to calculate the percentage of overlap (Mander’s coefficient) between Grl3p and V-ATPase subunits (Vma1p, Vma8p, Vma10p and V-ATPase-a1p) in cortical (I) and non-cortical compartments (J). The error bars show the SDs. (K-L) Co-expression of V-ATPase-a1p-2xmNeon and Grl3p-3xmCherry. In the surface section, there was complete overlap between Grl3p and V-ATPase-a1p, in the docked mucocysts. There was also significant co-localization of V-ATPase-a1p and Grl3p in non-cortical structures, as seen in cross-sections of the same cells. Scale bars, 10 μm.

The non-cortical puncta showed striking differences from those at the cortex. In contrast to the results for mature mucocysts, we found significant overlap between Grl3p and all V-ATPase subunits (Vma1p, Vma8p, Vma10p and V-ATPase-a1p) at cytoplasmic puncta that reflect biosynthetic mucocyst intermediates **(Figure 4D, 4F, 4H, 4J and 4L)**.

Taken together, these results argue that multiple V-ATPase subunits, known to form part of the enzymatically-active holo-V-ATPase complex, are present in immature mucocysts. However, only a subset of those subunits persist in mature mucocysts docked at the cell surface. These results suggest that disassembly of the V-ATPase complex occurs during mucocyst maturation. The incompleteness of the V-ATPase complex present in mature mucocysts provides an explanation for why these organelles do not maintain an acidic lumenal environment.

### *AP-3* is required for the de-acidification during mucocyst maturation

The data outlined above reveal that disassembly of the V-ATPase holo-complex occurs during the maturation process and does not depend on docking *per se* since it occurs in the MN173 mutant. To begin identifying requirements for disassembly, we examined genes known to be required for mucocyst maturation. *Tetrahymena* lacking the µ-subunit of the AP-3 trafficking adaptor do not synthesize mature mucocysts but instead accumulate large spherical cytoplasmic vesicles bearing the proGrls (Kaur et al., 2017). We expressed Grl3p-pHluorin-mCherry in Δ*apm3* and analysed co-localization using BIOP JACoP plugin **(Figure 5A-D)**. In Δ*apm3* cells, the mCherry-positive puncta predominantly lacked pHluorin signal, indicating that the lumen of these mucocyst intermediates is acidic **(Figure 5B-C, S4F and Movie S2)**. There were also puncta that showed green but not red signals, but these were consistently relatively few in number (**Figure 5D)**. It is known from studies in other organisms that deletion of AP-3µ leads to the loss of the entire AP-3 complex (Mullins et al., 2000; Cowles et al., 1997; Stepp et al., 1997). Our data therefore indicates that AP-3 activity is required for the transition from acidified mucocyst intermediates to neutral pH mature mucocysts.

**Figure 5.**
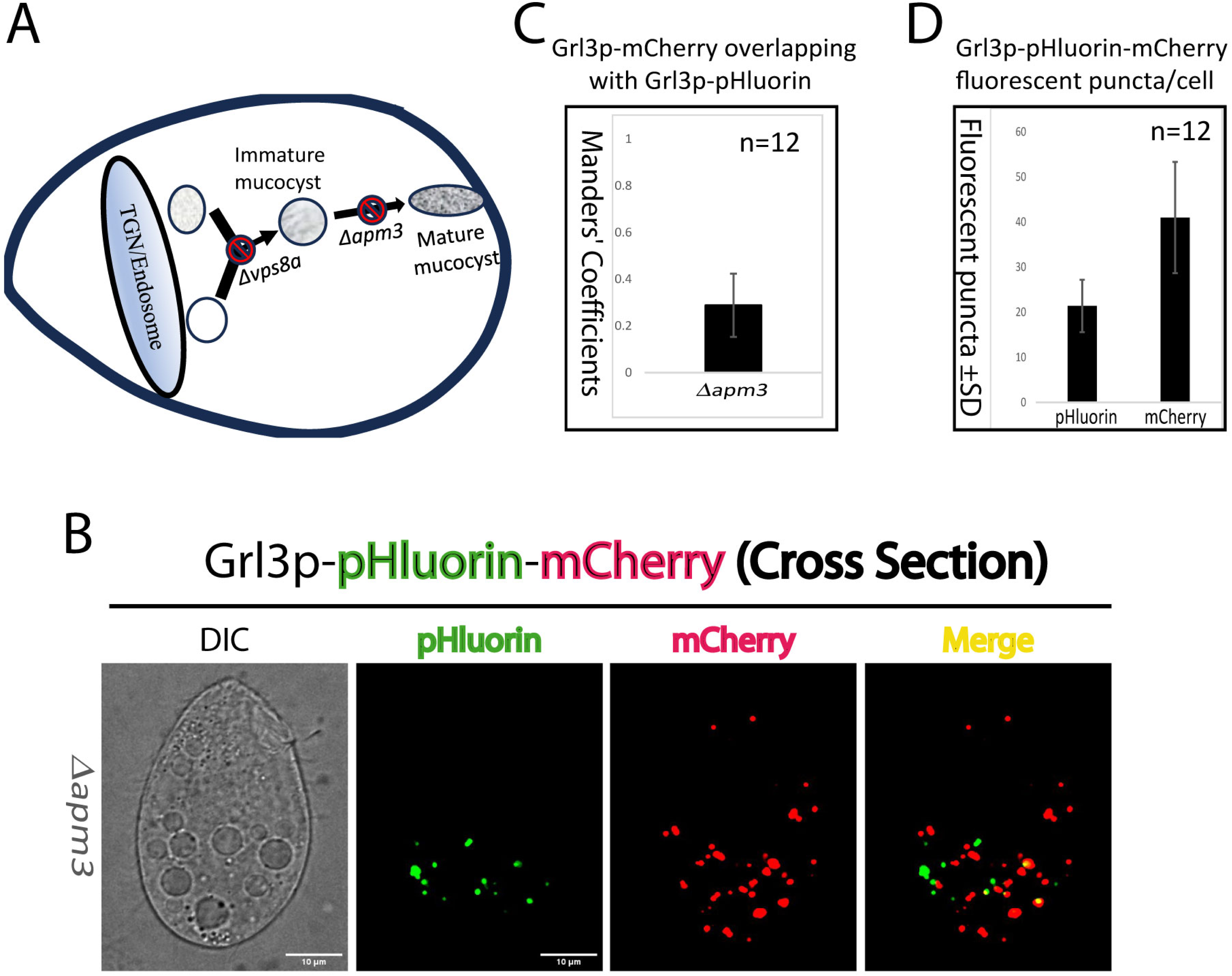
The disassembly of the V-ATPase holo-complex requires AP-3. (A) Cartoon depicting the maturation of mucocysts in *Tetrahymena*. (B) The co-localization of green and red signals from Grl3p-pHluorin-mCherry expressed in *Δapm3* cells, as shown in Figure 3E. In *Δapm3* cells, some co-localization of Grl3p was observed in the green and red channels within non-cortex compartments (C) Twelve separate images from panel B were analysed for non-cortical compartment. (D) Fluorescent puncta were quantified in both green and red channels from twelve separate images in panel B. The images were processed to remove noise, adjust thresholds and count fluorescent puncta using the Analyze Particles application in Image Fiji. Only particles ranging from 0.1 to 5 microns in size were counted to exclude background noise and large food vacuoles. The error bars show the SDs. Scale bars, 10 μm.

### *V-ATPase-a1* is required for secretion from mucocysts

To ask if *V-ATPase-a1* were required for mucocyst biogenesis, we subcloned the 5’UTR and 3’UTR of the *V-ATPase-a1* gene into a pNeo4 knockout vector and transformed it into *Tetrahymena* by biolistic transformation to target the *V-ATPase-a1* gene for disruption via homologous recombination with a drug-resistance cassette. (**Figure 6A)**. During ∼3-4 weeks of selection, all ∼45 copies of a gene in the polyploid macronucleus can be replaced with the cassette, resulting in a functional knockout if the gene is nonessential (Coyne et al., 2008). After generating the *V-ATPase-a1* knockout line, called *Δv-atpase-a1*, we used reverse transcription PCR (RT-PCR) to detect the gene transcript. The *V-ATPase-a1* transcript was not found in the deletion line (**Figure 6B**), which grew at wildtype rates under standard laboratory conditions. Hence the gene can be considered nonessential.

**Figure 6.**
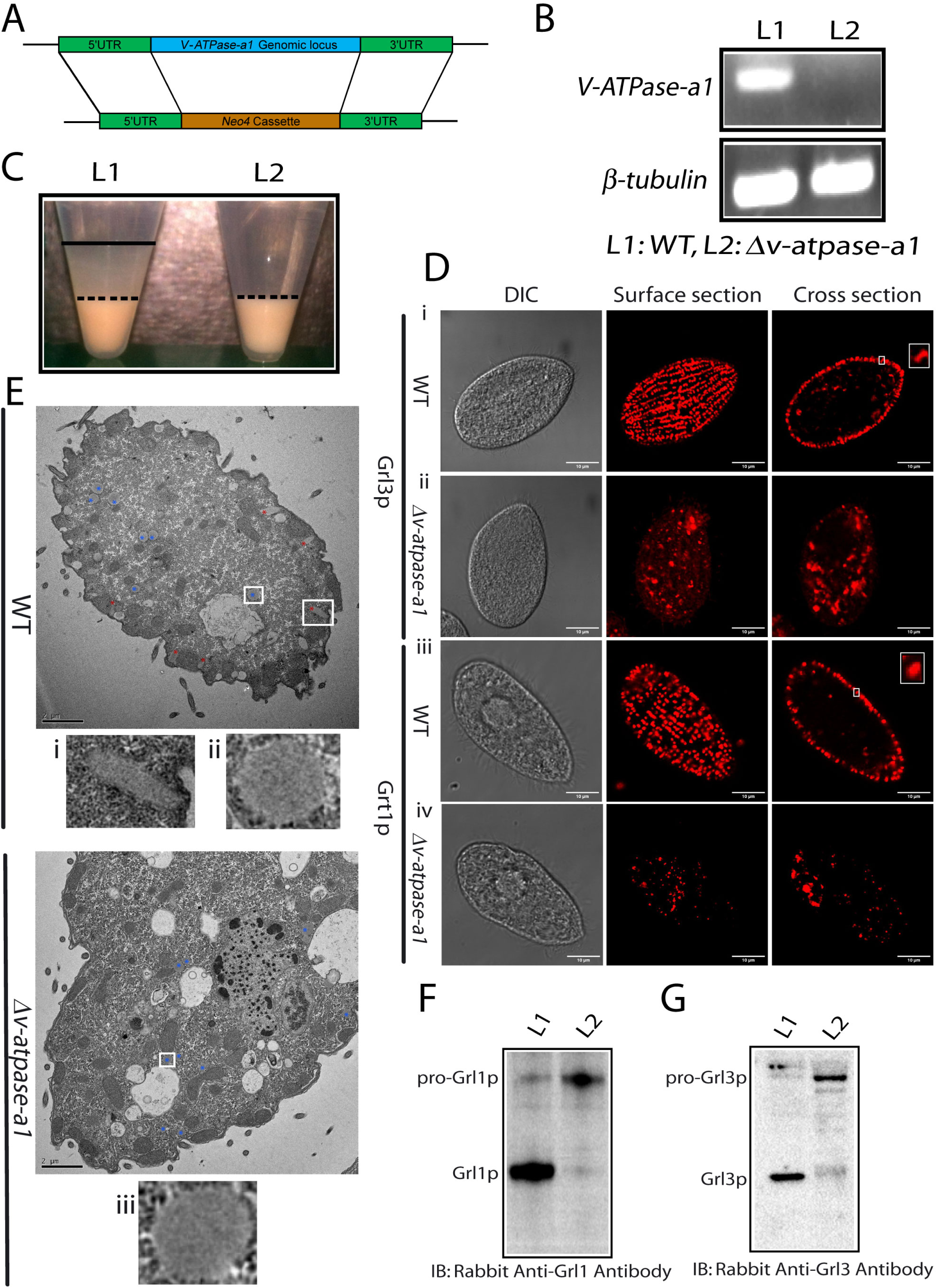
Analysis of the role of *V-ATPase-a1* in mucocyst biogenesis. (A) *V-ATPase-a1* knockout by homologous recombination. The knockout construct is described in Materials and Methods. (B) Confirmation of *V-ATPase-a1* knockout by one-step RT-PCR. PCR primers are listed in Supplemental **Table S3**. *BTU2* was used as a control. (C) *V-ATPase-a1* is required for the mucocyst secretion. Cells were stimulated with dibucaine to induce mucocyst exocytosis. Centrifugation results in a dense pellet of cells with an overlaying flocculent comprised of the secreted mucocyst cargo. The flocculent layer is delineated with a solid at the upper border and a dashed line at the lower border. The *Δv-atpase-a1* cells are grossly deficient in mucocyst secretion and produce no apparent flocculent layer (right) compared to the wildtype (left). (D-E) V-ATPase-a1p is essential for mucocyst formation. (D) Docked mucocysts visualized in fixed wild-type cells by anti-Grl3p mAb 5E9 (top panel i), and anti-Grt1p mAb 4D11 (lower panel iii). *Δv-atpase-a1* cells contain less mucocyst signal and the Grl3p puncta are often irregularly shaped (top ii and lower panel iv). Inset shows single docked mucocyst. Scale bars, 10 μm. (E) Electron micrographs show mucocysts in both wild-type (top) and *Δv-atpase-a1* (bottom). In wild-type, immature mucocysts (labelled with blue asterisks) and docked mature mucocysts (labelled with red asterisks) are present. In *Δv-atpase-a1*, no mature mucocysts are present. The inset displays a magnified view of the selected field. Scale bars, 2 μm. (F-G) V-ATPase-a1p is required for processing of proGrl proteins. Cell lysates (5000 cell equivalents in F and 10,000 cell equivalents in G) were separated by SDS-PAGE and Western blotted using antibodies against Grl1p and Grl3p. (F) Western blotting using rabbit anti-Grl1p antibody. The mature processed Gr1p product predominates in wild-type cells, while the unprocessed precursor of Grl1p predominates in *Δv-atpase-a1* cells. (G) Same as F but using anti-Grl3p antibody. Lane L1: WT and lane L2: *Δv-atpase-a1*.

Mucocysts are secretory organelles, so defects in biogenesis result in reduced cargo secretion. We therefore tested the exocytic response of the knockout cell line using an assay in which brief exposure of wild-type cells to the calcium ionophore dibucaine triggers simultaneous secretion from the several thousand mucocysts per cell (Satir, 1977; Cowan et al., 2005; Kumar et al., 2014).We found that *Δv-atpase-a1* cells were grossly deficient in mucocyst content release **(Figure 6C)**.

### *V-ATPase-a1* is essential for mucocyst formation

To test whether *V-ATPase-a1* was required for mucocyst formation *per se*, we used indirect immunofluorescence with two monoclonal antibodies (mAbs) that recognize, respectively, Grl3p (mAb 5E9) and Grt1p (mAb 4D11) (Cowan et al., 2005; Kumar et al., 2014). Imaging of wild-type cells using either antibody brightly illuminates the mucocysts primarily docked at the cell periphery, as seen in cell cross sections **(Figure 6Di, iii and inset)**. In contrast, in *Δv-atpase-a1* cells both Grl3p and Grt1p were relatively sparse, and no mucocysts were docked at the cortex (**Figure 6Dii and iv)**. Notably, large fractions of the Grl3p and Grt1p in the *Δv-atpase-a1* cells are distant from the cell cortex (**Figure 6Dii and iv)**. *V-ATPase-a1* therefore is required for mucocyst formation. To further confirm it, we examined wild-type and *Δv-atpase-a1* cells **(Figure 6E, S5)** by electron microscopy. In wild-type cells, both immature mucocysts, as well as mature mucocysts docked at the cell cortex, were detected **(Figure 6E, i and ii)**. In the *Δv-atpase-a1* mutant, only structures resembling immature mucocysts were observed **(Figure 6E, iii and S5**).

### *V-ATPase-a1* is required for the processing of pro-Grl proteins

The dense lumenal proteinaceous core of mucocysts in wild-type cells is composed of GRL proteins, and the formation of the mucocyst core depends upon their proteolytic processing (Cowan et al., 2005; Turkewitz et al., 1991). The morphologically aberrant mucocysts in *Δv-atpase-a1* cells may be caused by defective processing of Grl proteins **(Figure 6D-E**). Therefore, we asked whether *V-ATPase-a1* were required for Grl proprotein processing. We analysed whole-cell lysates of the *Δv-atpase-a1* cells by Western blotting using anti-Grl antisera **(Figure 6F-G)**. In the mutant cells, the Grl proteins were found to accumulate as unprocessed or partially-processed precursors **(Figure 6F-G)**.

### *V-ATPase-a1* is required for a key membrane trafficking step during mucocyst formation

In addition to proteolytic maturation of Grl proproteins, a well-documented step in mucocyst formation is heterotypic vesicle fusion that brings together the Grl proteins, which may be trafficked directly from the Golgi/TGN, with Grt proteins as well as at least some key proGRL processing enzymes which may be trafficked via endosomes. This fusion step involves a Syntaxin 7-like homolog, Stx7l1 and the CORVET subunit Vps8a (Sparvoli et al., 2018). Stx7l1p itself, but not Vps8ap, remains associated with mature mucocysts. In cells lacking either *VPS8a* or *STX7l1*, Grl and Grt proteins accumulate in separate vesicles.

To test whether V-ATPase-a1 is required for heterotypic vesicle fusion, we used indirect immunofluorescence to localize Grt1p, Grl1p and Stx7l1p in fixed cells. In wildtype cells, Grl1p and Grt1p accumulate exclusively in docked mature mucocysts **(Figure 7A, top panel)**. In contrast, in *Δv-atpase-a1* cells, both cargo proteins accumulate in a heterogeneous cohort of cytoplasmic vesicles **(Figure 7A, middle panel)**. Compared to that in wild-type cells, Grl1p and Grt1p show significantly less overlap (∼51%) in *Δv-atpase-a1* **(Figure 7B)**, which is similar to their overlap in *Δvps8a cells* **(Figure 7A, lower panel and 7B)**.

**Figure 7.**
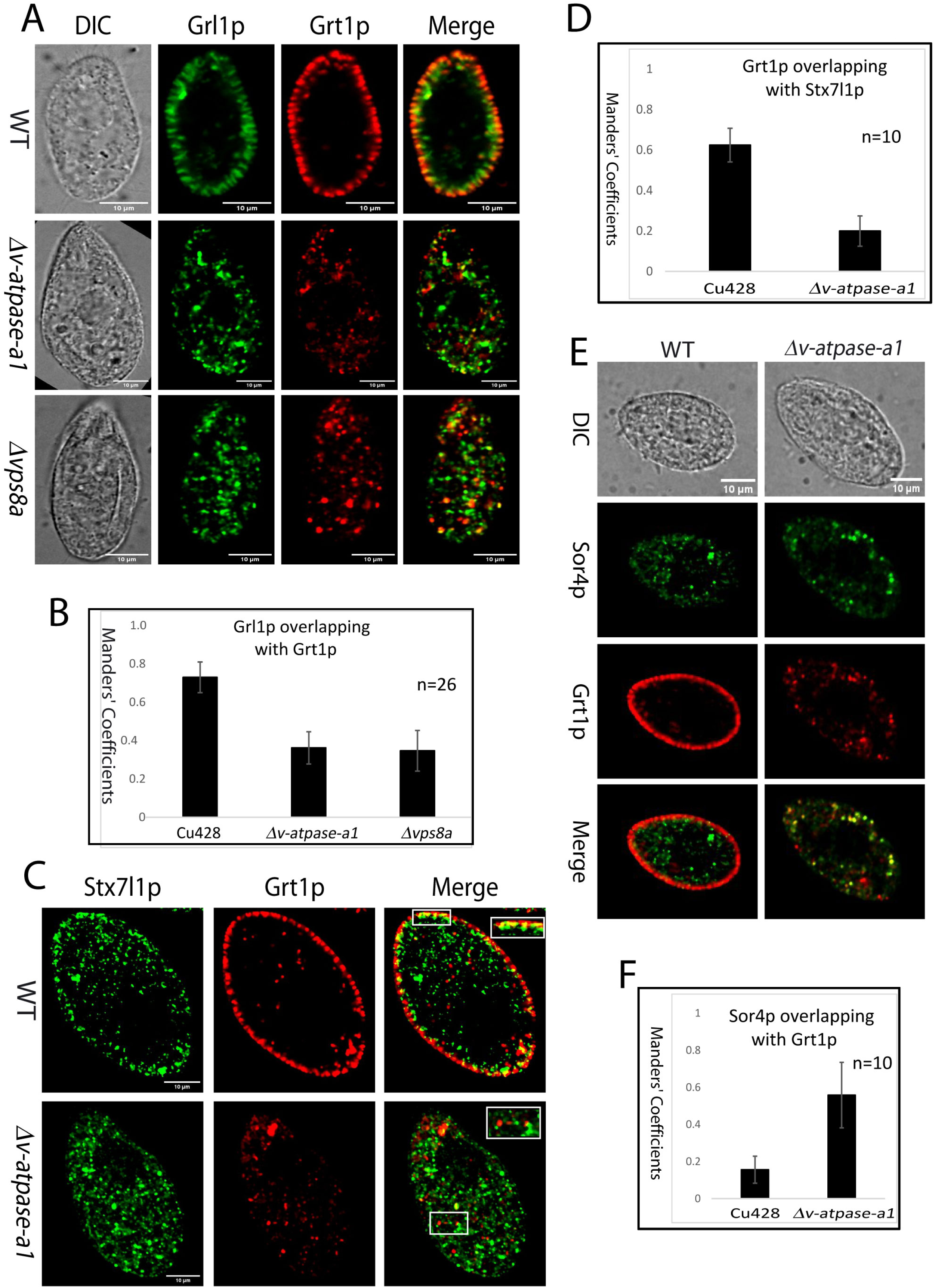
The *Δv-atpase-a1* cells show defects in vesicle trafficking during mucocyst biogenesis, similar to those in *Δvps8a* cells. (A) Wild-type (top panel), *Δv-atpase-a1* (middle panel) and *Δvps8a* (bottom panel) cells were double stained with rabbit anti-Grl1 and mouse monoclonal anti-Grt1 (4D11). Co-localization of Grl1p and Grt1p is significantly reduced in *Δv-atpase-a1* and *Δvps8a*, compared to wild-type. Optical cross sections are shown as single slices, for clarity. Scale bars, 10 μm. (B) Percentage of overlap (Mander’s coefficient) between Grl1p and Grt1p using a sample of 26 non-overlapping images. The error bars show the SDs. (C) Wild-type (top panel) and *Δv-atpase-a1* (bottom panel) expressing Stx7l1p-GFP were double stained with rabbit anti-GFP and mouse monoclonal anti-Grt1(4D11) antibodies. In *Δv-atpase-a1* cells, there is a significant decrease in the co-localization of Stx7l1p and Grt1p. Inset shows co-localization between Stx7l1p and Grt1p. (D) A sample of ten non-overlapping fields was analyzed to determine the percentage of overlap (Mander’s coefficient) between Stx7l1p and Grt1p using the Fiji-JACoP plugin. (E) Wild-type (left panel) and *Δv-atpase-a1* (right panel) cells expressing Sor4p-GFP were double stained with rabbit anti-GFP and mouse monoclonal anti-Grt1 antibodies. (F) The Fiji-JACoP plugin was used to calculate the percentage of overlap (Mander’s coefficient) between Sor4p and Grt1p as before. Scale bars, 10 μm. The difference in cell size between samples is due to variable flattening by the coverslips.

Consistent with previous results in wildtype cells (Kaur et al., 2017), Stx7l1p-GFP co-localized with Grt1p in docked mucocysts **(Figure 7C, top panel and S6A)**. Their co-localization is ∼68% reduced in *Δv-atpase-a1* cells **(Figure 7C-D)**. Notably, a greater extent of co-localization is observed in the mutant cells between Grl3p and Stx7l1p **(Figure S6B-C)**. These results are consistent with earlier reports that Stx7l1p is associated with vesicles containing Grl3p rather than Grt1p (Kaur et al., 2017; Sparvoli et al., 2018). Our results suggest that the absence of V-ATPase-a1 inhibits heterotypic fusion between Grl-and Grt-containing vesicles.

We previously found evidence that the targeting of Grt1p to mucocysts depends on its interaction with a VPS10-family receptor, Sor4p (Briguglio et al., 2013). The Grt1p-containing vesicles in *Δv-atpase-a1* cells, if they are stalled trafficking intermediates, might therefore also contain Sor4p. In agreement with this model, we found that the co-localization of Grt1p and Sor4p increased 3.5-fold in the *Δv-atpase-a1* mutant compared to in wildtype cells **(Figure 7E-F, S6D)**. Taken together, our results support the idea that V-ATPase-a1p promotes fusion of Grl1p-containing and Sor4p/Grt1-containing vesicles during mucocyst formation, showing phenotypes similar to those seen in *Δvps8a* cells. Based on these data, we proposed a model for the mucocyst biogenesis pathway in *T. thermophila* **(Figure 8)**, extending the model proposed by Sparvoli *et al,* 2018.

**Figure 8.**
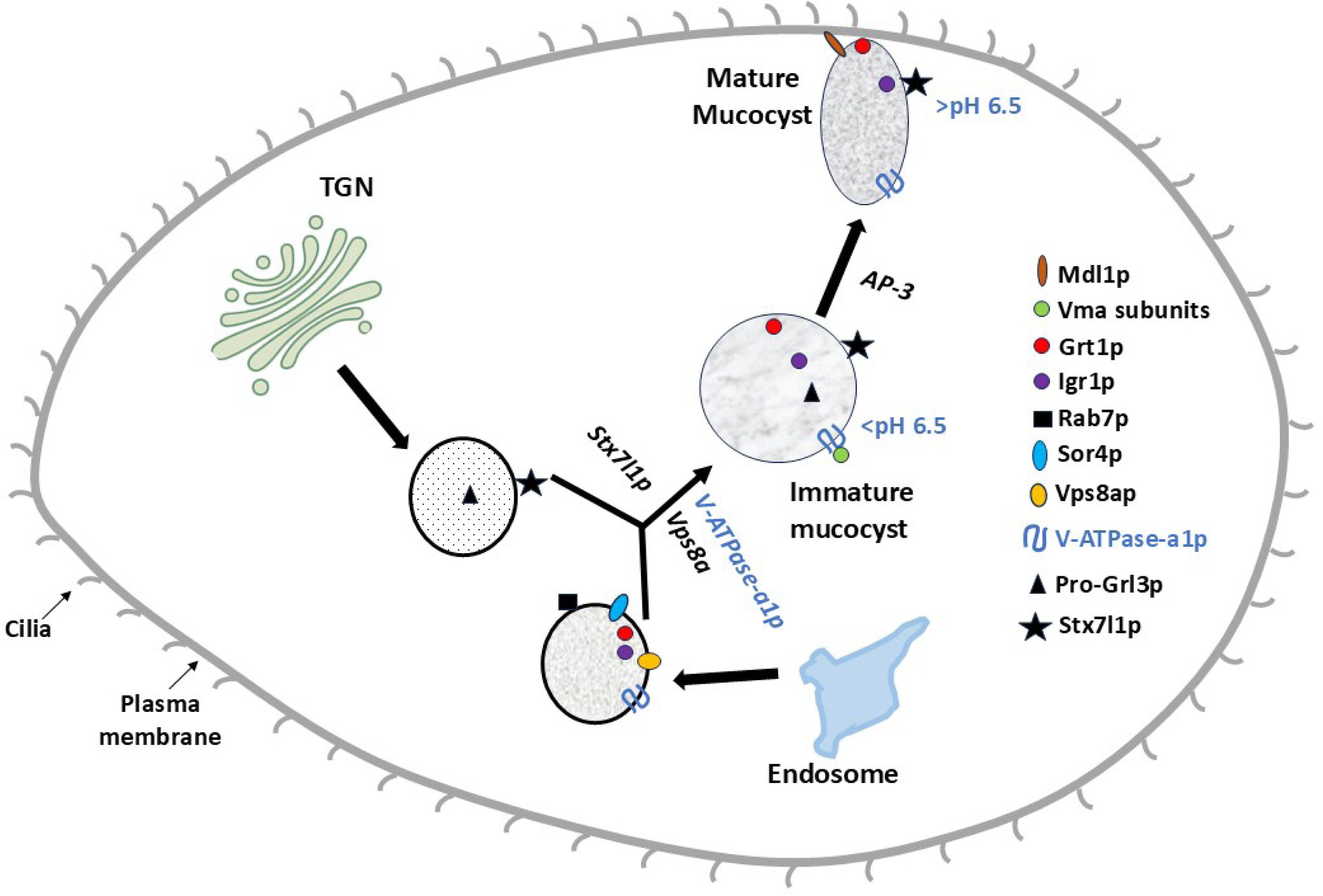
Proposed model for the mucocyst biogenesis pathway in *T. thermophila*, extending that proposed by Sparvoli et al., 2018. Vps8ap and Stx7l1p are required for homotypic and/or heterotypic fusion of vesicles bearing pro-Grls and other mucocyst cargo proteins, to generate immature mucocysts whose subsequent maturation depends on processing enzymes and the AP-3 adaptor. We now include V-ATPase-a1 in this model, which appears to be required for the same step of efficient vesicle fusion to form immature mucocysts. These are transiently acidified, and subsequently Vma subunits are lost via V-ATPase complex disassembly during mucocyst maturation. At the end of this process, the mature docked mucocysts are non-acidic. The disassembly of the V-ATPase holo-complex appears dependent on AP-3, since the immature mucocysts that accumulate in *Δapm3* are acidified.

## Discussion

In *Tetrahymena thermophila*, the biogenesis of secretory vesicles known as mucocysts has previously been shown to depend on a set of proteins associated in other lineages with LROs (Kaur et al., 2017; Sparvoli et al., 2018). Here, we investigated whether a V-ATPase was also required for mucocyst formation. Based on genome annotation, *T. thermophila* expresses a large number of distinct V-ATPase subunits, many of which are likely to be compartment-specific. Among the 6 *T. thermophila* paralogs encoding a-subunits of the V_0_ subcomplex, we discovered that the a1 paralog is transcriptionally coregulated with known mucocyst-associated genes, and that the corresponding protein localizes strongly, though not exclusively, to mature docked mucocysts. In other organisms, the localization specificity of V-ATPases has been shown to depend on determinants in the a-subunit of the V_0_ subcomplex (Sun-Wada and Wada, 2013, 2022; Sun-Wada et al., 2007; Matsumoto et al., 2018; Oka et al., 2001; Chen et al., 2024; Capecci and Forgac, 2013; Toei et al., 2010). Consistent with this idea, in animal cells distinct a-subunits are found respectively in coated vesicles, the Golgi apparatus/early endosomes, and late endosomes/lysosomes (Breton and Brown, 2013; Chu et al., 2021; Matsumoto et al., 2018; Toei et al., 2010). The six genes encoding a-subunits of the V-ATPase in *T. thermophila* compare to just four a-subunit genes in mammals (Sun-Wada and Wada, 2013; Toei et al., 2010) while 17 a-subunit genes are present in *Paramecium tetraurelia* (Wassmer et al., 2006).

In *Paramecium,* these seventeen genes fell into nine families and localized to at least seven different compartments. Notwithstanding the localization of a V-ATPase subunit to mucocysts, we demonstrate that they are similar to neutral pH storage LROs that have been documented in mammals, Amoebozoa, and other lineages. That is, we found by three different approaches that mature mucocysts, which dock at the plasma membrane, are not detectibly acidic. The most impactful of these approaches was based on tagging a mucocyst-cargo protein with pHluorin, a fluorophore whose emission is pH-sensitive, because it also allowed us to detect and analyze immature mucocysts. Using cells expressing this construct, we could detect mucocyst intermediates, i.e., cytoplasmic vesicles containing the mucocyst cargo protein Grl3p, that were acidified in contrast with mature mucocysts. Thus, our data strongly suggest that mucocysts are transiently acidified during their biogenesis. We found that the a1 subunit gene is non-essential but required for mucocyst biogenesis. Similarly, the a3 subunit in *Paramecium* had been implicated in the formation of trichocysts, which are homologous to mucocysts (Wassmer et al., 2006). The transient acidification of ciliate secretory LROs at an intermediate phase in organellar maturation may be sufficient to explain the defects seen with knockout or silencing of the V-ATPase complex subunit genes. At the same time, it remains possible that specific subunits, in either the intact or partially disassembled complex, play roles distinct from proton pumping.

In *Tetrahymena* we detected numerous defects in mucocyst formation in a1-knockout cells, including failure to process Grl proproteins as well as likely defects in heterotypic vesicle fusion that is a normal feature of mucocyst biogenesis. The proprotein processing defects are similar to those we previously found in cells lacking the gene encoding the key processing enzyme in mucocysts, *CTH3* (Kumar et al., 2014). These similarity in defects can be readily explained if transient acidification of the immature mucocyst is required to activate Cth3p. Similarly, the apparent defects in vesicle fusion, which resemble those in cells lacking the Vps8a subunit of a mucocyst-specific CORVET complex, could be indirect consequences of a failure to acidify one or more intermediates. However, such defects could also reflect a more direct role of one or more V-ATPase complex subunits in this step. The possibility that V-ATPases participate in some membrane fusion reactions, via mechanisms unrelated to their proton-pumping activity, has been a contentious issue, but includes recent support for a direct fusogenic role (Sreelatha et al., 2015; Strasser et al., 2011; Coonrod et al., 2013; El Far and Seagar, 2011; Wang and Hiesinger, 2013). Interestingly, in mammalian cells there is evidence that the V-ATPase on secretory vesicles acts to acidify the lumen but also to regulate exocytosis machinery by acting as a pH sensor (Poëa-Guyon et al., 2013). The issue of potential subunit activities that are independent of proton pumping is also relevant for subunits like a1 that remain mucocyst-associated after the holo-complex has disassembled during maturation.

We found that a significant fraction of a1-positive cytoplasmic puncta are also positive for Rab7. Rab7 was previously shown to be present on vesicles containing the mucocyst-destined protein Grt1p (Sparvoli et al., 2018). Consistent with this, we found significant colocalization between V-ATPase-a1p and Grt1p in cytoplasmic puncta that are likely intermediates in mucocyst biogenesis. The association of V-ATPase with Rab7 has also been reported in other cell types (Matsumoto et al., 2022, 2018; Matsumoto and Nakanishi-Matsui, 2019; Nakanishi-Matsui et al., 2024). Studies in osteoclasts demonstrated that subunit a3 recruits Rab7 to secretory lysosomes, promoting their transport to the cell periphery where they deliver bone-resorbing enzymes together with the proton pump. The a3 subunit has also been shown to be involved in the transport of Mon1A-Ccz1, a guanine nucleotide exchange factor (GEF) for Rab7, to secretory lysosomes (Matsumoto et al., 2022). Thus, the non-enzymatic roles of V-ATPase subunits include determinants for organelle transport (Matsumoto et al., 2018, 2022). The physiological regulation of V-ATPase in mammals involves its reversible assembly/disassembly, among other mechanisms. For example, in yeast and mammalian cells such reversible assembly of the V_1_ and V_0_ domains regulates endolysosomal/lysosomal pH (Lafourcade et al., 2008; Sava et al., 2024). Our results suggest that a similar phenomenon underlies the properties of mucocysts in *Tetrahymena*, unfolding in the context of an organellar maturation pathway. We found that V_1_ subunits Vma1p, Vma8p and Vma10p are not present in mature, docked mucocysts, whereas they instead are found in a pool of cytoplasmic Grl3-positive puncta that are likely to represent immature mucocysts. These results suggest that the mucocyst-associated V-ATPase complex partially disassembles during maturation, with only a subset of the subunits remaining mucocyst-associated, which is correspondingly non-acidic. Importantly, at least one of the dissociated subunits, Vma8, has previously been shown to be essential for coupling of proton transport and ATP hydrolysis in yeast (Graham et al., 1995; Xu and Forgac, 2000). Therefore, the remnant V-ATPase complex in mature mucocysts is likely to be disabled for proton translocation, which can explain why mucocysts do not maintain an acidic luminal environment. Because non-acidic storage LROs are not unique to ciliates, similar phenomena could be an important feature of LRO specialization in other lineages.

The heterotetrameric trafficking adapter AP-3 is considered a conserved factor in LRO formation, and more broadly has a role in mediating cargo transport from the late/trans-Golgi to late endosomal compartments (Leih et al., 2024; Ma et al., 2021). We previously found that the AP-3 adapter was required for mucocyst formation, and speculated that it delivered an essential maturation factor. While the work in our current manuscript does not advance our understanding of the direct interactors of AP-3 in this pathway, we can now add significantly to the model by showing that AP-3 activity is required to promote disassembly of V-ATPase during mucocyst maturation. On this basis, we propose that AP-3 be considered a regulator of V-ATPase activity.

## Materials And Methods

### Reagents, *Tetrahymena* Strains and culture conditions

*Tetrahymena thermophila* strains were grown overnight at 30°C with agitation in nutrient rich SPP medium (2% proteose peptone, 0.2% dextrose, 0.1% yeast extract, 0.003% ferric EDTA) supplemented with 250 μg/ml streptomycin sulfate, 250 μg/ml penicillin G, and 0.25 μg/ml amphotericin B fungizone, to medium density (2-3 × 10^5^ cells/ml). Cu428 wild type strain was used as a control for all experiments and for biolistic transformation. 10mM Tris buffer (pH 7.4) was used for washing cells and as a starvation medium to reduce autofluorescence in food vacuoles. Cell culture densities were measured using a hemocytometer. *T. thermophila* strains used in this study are listed in **Table 1**. All reagents, chemicals, plasmids, and other information are listed in Key Resources Supplemental **Table S2**. The constructs p2xHA-3xmCherry-Rab7p-NCVB, pGrl3p-GFP-CHX, pSor4p-GFP-CHX, p-2xmNeon-6c-myc-Neo4, p3xmCherry-2xHA-Neo4 and *Δvps8a*-Neo4 were from (Sparvoli et al., 2018); pmEGFP-Neo4 and pNeo4 KO vector were from (Kumar et al., 2014); pStx7l1p-GFP-Blasticidin was from (Kaur et al., 2017). Plasmids pPOC1-mCherry-BSR and pPur4 were generously provided by Prof. Chad Pearson (University of Colorado, USA) and Dr. Kazufumi Mochizuki (University of Montpellier, France), respectively. Wild type strain Cu428 was from Dr Abdur Rahaman (NISER, India).

**Table 1:**
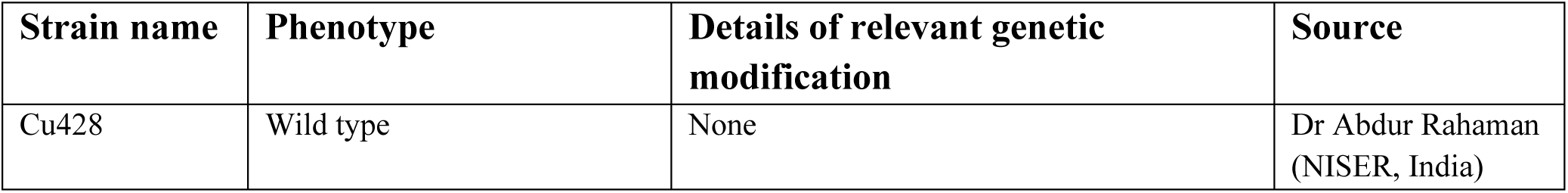
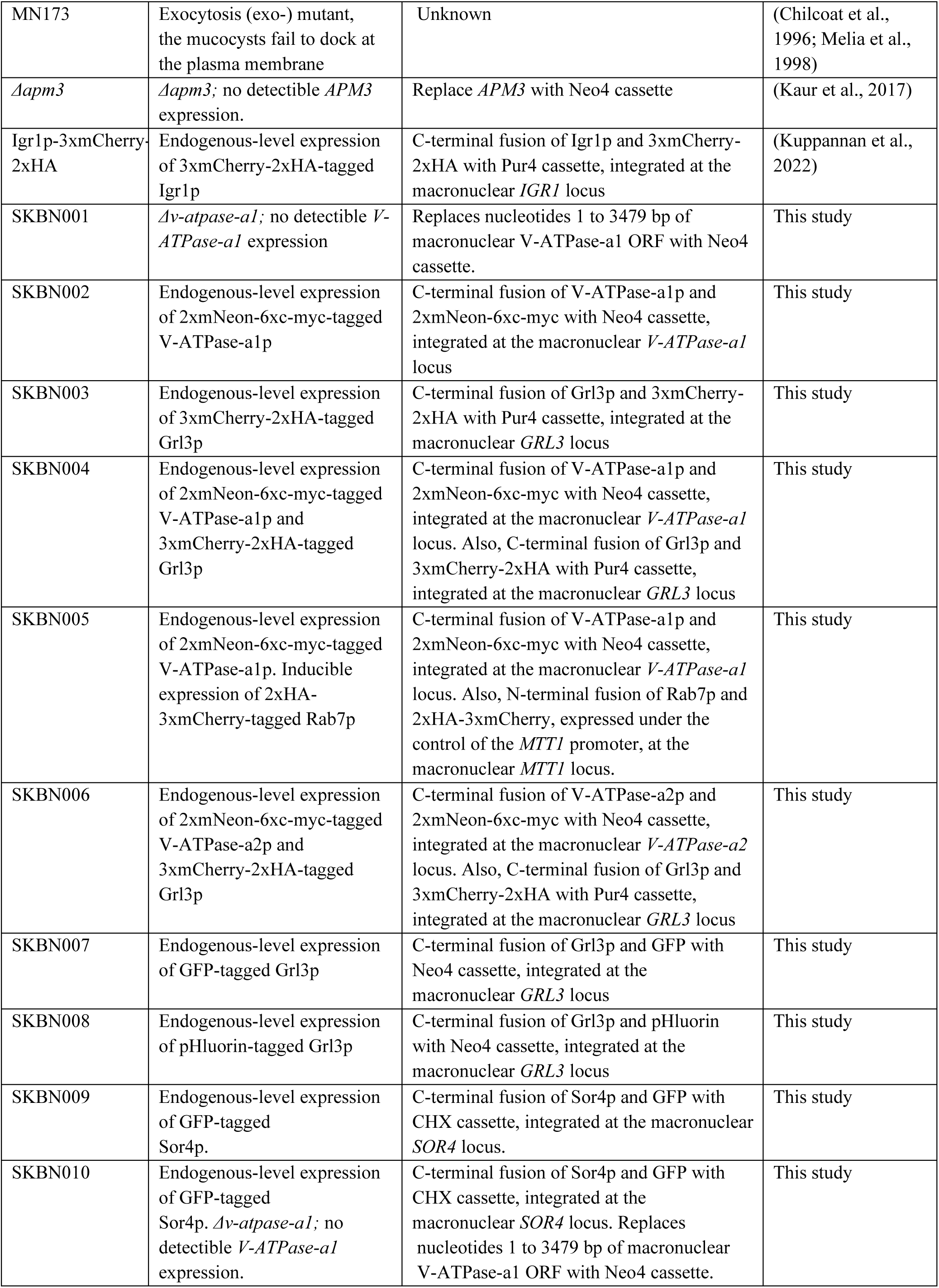
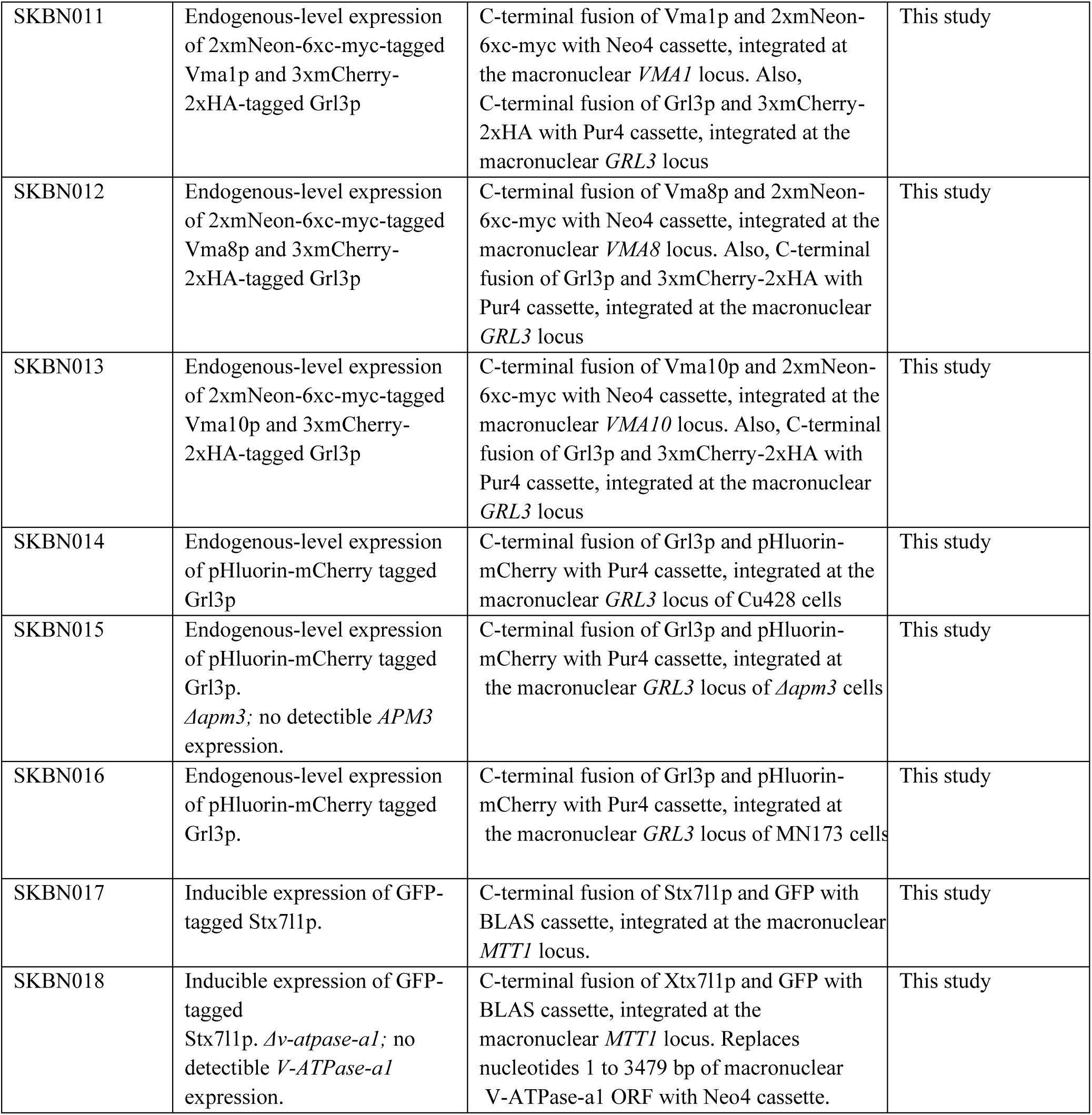
*Tetrahymena* Strains used in this study.

### Exocytosis assay

Exocytosis was stimulated by dibucaine as described previously (Kumar et al., 2014; Rahaman et al., 2009).

### Verification of knockouts of genes by reverse transcriptase (RT)-PCR

RT-PCR was performed as described previously (Kumar et al., 2014). Briefly, total RNA was extracted by following the manufacturer’s instructions using RNeasy mini spin column kits (Qiagen, USA). cDNA (200-500 base pairs of each gene) was synthesized and further amplified using the one step RT-PCR kit (Qiagen USA,) as per manufacturer’s protocol using primers designated in **Supplemental Table S3**. To provide positive controls, RT-PCR using β-tubulin specific primers were performed in parallel.

### Biolistic transformation

Biolistic transformations were carried out as described (Kumar et al., 2015, 2014) with some modifications. Briefly, cells were grown in SPP medium and incubated at 30^°^C with swirling to reach medium cell density. The next day, cells were washed twice with 10 mM Tris pH 7.4 and starved in the same buffer for 16 to 18 hours. 15 μg of total linearized plasmid DNA were conjugated with 0.6 μ gold particles (Bio-Rad, USA) as per the manufacturer’s instructions. Four hours after bombardment, drugs were added to cultures that were swirling at 30°C to select positive transformants. Positive transformants were identified after 3–4 days and serially transferred 5 days/week in increasing concentrations of drug and decreasing concentrations of CdCl_2_ (up to 6.5 mg/ml of paromomycin, 0.1 μg/ml CdCl_2_; up to 200 μg/ml of puromycin and 0.5 μg/ml CdCl_2_; up to 60 μg/ml of blasticidin and 0.5 μg/ml CdCl_2_, 20 μg/ml of cycloheximide and 0.8 μg/ml CdCl_2_) for 4-5 weeks before further testing. Each line was evaluated using at least three independent transformants.

### Generation of *V-ATPase-a1* knockout strain

Upstream (∼1 kb) and downstream (∼1 kb) regions of the *V-ATPase-a1* (TTHERM_01332070) gene were amplified by PCR and subcloned into the SacI and XhoI sites of the pNeo4 vector, respectively, using the In-Fusion cloning kit (TaKaRa, Japan). *V-ATPase-a1* knockout construct was linearized using KpnI and SapI enzymes before biolistic transformation into Cu428 cells. The entire sequence of the *V-ATPase-a1* gene was deleted. The primers used to amplify target regions are listed in **Supplemental Table S3**.

### Live-cell microscopy

Cultures for live microscopy were analyzed at mid-density unless otherwise indicated. To image cells expressing mNeon, mCherry, GFP or pHluorin-tagged fusion proteins, transformants were cultured overnight in SPP media and then transferred into 10 mM Tris pH 7.4 for 2-4 h to reduce autofluorescence in food vacuoles. Co-localization in live cells was determined by simultaneous imaging of mNeon or GFP-tagged fusion proteins with mCherry-tagged mucocyst markers. To simultaneously localize mNeon tagged V-ATPase-a1p with Lyso-Tracker Red (Invitrogen, USA), cells were incubated with 200 nM Lyso-Tracker Red for 5 min and images were captured as previously described (Kumar et al., 2014). For co-localization of V-ATPase-a1p and FM4-64 (Invitrogen, USA), cells were treated with 5 μM FM4-64, which labels endosomes, for 5 minutes and then pelleted and resuspended in 10 mM Tris pH 7.4. Subsequently, cells were imaged within 30-40 minutes. Cells expressing Igr1p-3xmCherry were incubated with 0.1 µg/ml Acridine Orange (Invitrogen, USA) for 2 min, while cells expressing Grl3p-3xmCherry were incubated with 5 µM Protonex Green (AAT Bioquest, USA) for 120 min, both at room temperature and in the dark. Cells were washed twice with 10 mM Tris pH 7.4 and observed under the microscope. To check co-localization between V-ATPase-a1p and Rab7 positive endosomes, cells were transformed to co-express V-ATPAse-a1p at the endogenous locus with mCherry-Rab7p. Transgene mCherry-Rab7p was induced for 2h in SPP with 2 μg/ml CdCl_2_ followed by induction with 0.5μg/ml CdCl_2_ for 2h in starvation buffer 10 mM Tris, pH 7.4. Stx7l1p-GFP was induced with CdCl_2_ as mentioned above. Live-cell images were captured using an Olympus SR10 spinning disk super-resolution confocal microscope (Olympus, Japan). Both confocal and super-resolution modes with a 60x oil objective were used to capture time-lapse videos. For z sectioning, SORA disk-enabled acquisition was used to keep the z slice step size at 0.3 μm. After removing the background signal from the images, they were saved as JPEGs that were denoised, colored, and adjusted for brightness and contrast using the Fiji application (http://fiji.sc/Fiji).

### Immunofluorescence Assay

Cells were grown to reach a cell density of 5×10^5^ cells/ml, then starved in 10 mM Tris pH 7.4 for 2 hours, fixed, and immunolabeled as described previously (Kumar et al., 2014). Monoclonal antibodies 5E9 [1:9; (Bowman and Turkewitz, 2001)and 4D11 1:5; (Turkewitz and Kelly, 1992)] were used to visualize Grl3p and Grt1p, respectively followed by goat anti-mouse conjugated with Texas red (1:100; Invitrogen, USA). Grl1p and GFP-tagged fusion proteins were visualized by 1:5000 diluted rabbit anti-Grl1 (Kuppannan et al., 2022; Ota, 2018), and rabbit anti-GFP (1:400; (Invitrogen, USA), respectively, followed by anti-rabbit antibody conjugated with Alexa 488 (1:250 Invitrogen, USA). Cells were double immunolabeled with mouse mAb 4D11 and rabbit anti-Grl1 to visualize the fusion of Grl1p and Grt1p-containing vesicles. Fixed cells were imaged on a Leica SP5 II STED-CW super-resolution laser scanning confocal microscope (Leica, Germany) or an Olympus SR10 spinning disk super-resolution confocal microscope. Images were analyzed as described above.

### Electron microscopy

Cells were grown to reach high cell density (10^6^/ml), washed with 10mM Tris media (pH 7.4), fixed, section stained and imaged as described previously (Kumar et al., 2015).

### SDS-PAGE, and Western blotting

Whole cell lysates were prepared from 3×10^5^ cells as described (Kumar et al., 2014). Using mouse anti-c-Myc Agarose beads (Thermo Fisher Scientific, USA), 2xmNeon-6c-myc-tagged fusion proteins were immunoprecipitated from detergent lysates as previously described (Sparvoli et al., 2018). Samples were separated using SDS-PAGE and then transferred to 0.45μ PVDF membranes (Merck Millipore, USA) for western blots. Blots were blocked and probed with antibodies described previously (Turkewitz et al., 1991). The primary antibodies rabbit anti-Grl1, rabbit anti-Grl3 (Kumar et al., 2014, 2015), mouse monoclonal anti-GFP (Biolegend, USA) and anti-c-Myc (Sigma, USA) were diluted 1:2000, 1:800, 1:5000 and 1:4000, respectively. Proteins were visualized using Super Signal West Femto Maximum Sensitivity Substrate (Thermo Fisher Scientific, USA), and enhanced chemiluminescence horseradish peroxidase–linked anti-rabbit (Bio-Rad, USA), anti-mouse (Bio-Rad, USA),) and anti-mouse for IP (Abcam, UK), with the secondary antibodies diluted 1:5000, 1:5000, and 1:1000, respectively.

### Expression of V-ATPase-a1p, V-ATPase-a2p, Vma1p, Vma8p, Vma10p, Cth3p, Grl3p and Igr1p at endogenous locus

The p2xmNeon-6c-myc-Neo4 and pmEGFP-Neo4 vectors were previously described (Sparvoli et al., 2018; Kumar et al., 2014; Briguglio et al., 2013). To create pV-ATPase-a1p-2xmNeon-6c-myc-Neo4, the genomic DNA of V-*ATPase*-a1 (TTHERM_01332070; 1462bp) without stop codon and 1000bp of V-*ATPase*-a1 downstream genomic region were amplified and cloned into p2xmNeon-6c-myc-Neo4 vector at NotI and XhoI sites, respectively, using In-Fusion Cloning (TaKaRa, Japan). The construct was named pV-ATPase-a1p-2xmNeon-6c-myc-Neo4 was linearised with KpnI and SacI, 2xmNeon-6c-myc was fused to the C-terminus of V-*ATPase*-a1 at the endogenous macronuclear locus via homologous recombination. The similar approach was used to amplify the genomic DNA without stop codon (∼700-900bp) and downstream genomic sequence (∼600-850bp) of other V-ATPase subunits *V-ATPase-a2* (TTHERM_00463420), *VMA1* (TTHERM_00339640), *VMA8* (TTHERM_00821870) and *VMA10* (TTHERM_00052460) for fusion of 2xmNeon-6c-myc to the C-terminus of V-ATPase subunits at the endogenous macronuclear locus. For the generation of pGrl3p-3xmCherry-2xHA-Pur4, the pGrl3p-GFP-CHX vector was used as template (Sparvoli et al., 2018). BamHI/SpeI sites were used to replace monomeric mGFP with p3xmCherry-2xHA, while PstI/HindIII sites were used to replace CHX (cycloheximide resistance gene) with Pur4 (puromycin resistance gene) to generate pGrl3p-3xmCherry-2xHA-Pur4 vector. The pmEGFP-Neo4 and pGrl3p-3xmCherry-2xHA-Pur4 vectors were used to make the pGrl3p-mEGFP-Neo4 construct. The genomic (857 bp excluding stop codon) and downstream sequence (783bp) of *GRL3* gene were excised from pGrl3p-3xmCherry-Pur4 vector using BamHI/ SacI and HindIII/KpnI, respectively and ligated into pmEGFP-Neo4 vector at mentioned sites respectively using T4 DNA ligase (New England, Biolabs, USA). pGrl3p-mEGFP-Neo4 vector was used as template to generate pGrl3p-pHluorin-Neo4 construct. Super ecliptic pHluorin (SEP) variant sequence was derived from the pHluorin MT1-MMP vector (Lizárraga et al., 2009). The pHluorin gene (lacking stop codon), codon optimised for *Tetrahymena* using the IDT Codon optimization tool, was synthesized (Eurofins Genomics Pvt Ltd) with added BamHI/SpeI sites at 5’ and 3’ end of gene. Monomeric mGFP from pGrl3p-mEGFP-Neo4 was replaced with pHluorin using BamHI/SpeI sites to generate pGrl3p-pHluorin-Neo4. To construct pGrl3-pHluorin-mCherry-Pur4, the Neo4 cassette of pGrl3p-pHluorin-Neo4 was replaced by Pur4 from pGrl3-3xmCherry-Pur4 using Kpn1 and Pst1 sites to generate pGrl3-pHluorin-Pur4. mCherry gene fragment was amplified from pPOC1-mCherry-BSR using primers designated in **Supplemental Table S3** and cloned into pGrl3-pHluorin-Pur4 vector at the SpeI site using In-fusion cloning. All constructs were confirmed by DNA sequencing. The final constructs were linearized with SacI and KpnI before transformation. The p2xHA-3xmCherry-Rab7p-Ncvb construct was linearized by digestion with SfiI and transformed into cells expressing V-ATPase-a1p by biolistic transformation. pGrl3p-3xmCherry-2xHA-Pur4, pGrl3p-mEGFP-Neo4 and pGrl3p-pHluorin-Neo4 were transformed into wildtype Cu428 using biolistic transformation. pGrl3-pHluorin-mCherry-Pur4 vector was transformed into Cu428, *Δapm3* and MN173 cells. V-ATPase-a1p-2xmNeon-6c-myc-Neo4, pV-ATPase-a2p-2xmNeon-6c-myc-Neo4, pVma1p-2xmNeon-6c-myc-Neo4, pVma8p-2xmNeon-6c-myc-Neo4 and pVma10p-2xmNeon-6c-myc-Neo4 vectors were transformed into Cu428 cell expressing Grl3p-3xmCherry-2xHA-Pur4.

### Phylogenetic tree construction

Protein BLAST (BLASTp) was used to identify homologs of V-ATPase-a1p (TTHERM_01332070; XP_001010002.1) in ciliates (*Paramecium*), apicomplexans (*Toxoplasma gondii), Arabidopsis thaliana, Caenorhabditis elegans, Drosophila melanogaster, Danio rerio,* and *Homo sapiens*, as shown in Supplemental **Table S4**. For tree construction, the top hits from each lineage were selected, assembled, and aligned using ClustalX (1.8). Phylogenetic trees were built using MEGA 11 software (Molecular Evolutionary Genetics Analysis: www.megasoftware.net/) with the maximum likelihood option, without gap regions and with1000 bootstrap replicates.

### In silico analyses

Protein sequences of V-ATPase-a subunit paralogs (a1p-a6p) were extracted from TGD (https://tet.ciliate.org/) and aligned using EMBOSS Needle (https://www.ebi.ac.uk/jdispatcher/psa/emboss_needle) using the default parameters. Transcription profiles were downloaded from the Tetrahymena Functional Genomics Database (http://tfgd.ihb.ac.cn) (Xiong et al., 2013, 2011; Miao et al., 2009). For plotting, each profile was normalized by setting the maximum expression level of the gene to 1. The V-ATPase-a1p structure was predicted using Protein Homology/Analogy Recognition Engine V 2.0 (Phyre 2), a web portal for protein modelling (Kelley et al., 2015). Structural homologs of the *T. thermophila* V-ATPase-a1p were identified using the AlphaFold Protein Structure Database (https://alphafold.ebi.ac.uk/).

## Abbreviation

LRO: Lysosome-related organelle
DCG: Dense core granule
GFP: Green fluorescent protein
V-ATPase: Vacuolar-ATPase
SEP: Super-ecliptic pHluorin
Igr1: Induced during granule regeneration 1
Vps8: Vacuolar protein sorting 8
Stx7l1: Syntaxin 7-like 1
Grl: Granule lattice
Grt: Granule tip
AP-3: Adaptor protein complex-3
VMA: Vacuolar Membrane Atpase

## Acknowledgements

We thank Dr Prasad Abnave (NCCS, Pune) for critical evaluation and valuable feedback on the manuscript and Dr Jomon Joseph for many helpful discussions. We heartily thank our lab members Shubham Yadav and Satya Santoshi for their continuous and generous support; Nikhat Khan for skilful technical assistance in all the above experiments. We are grateful to the confocal microscopy facility of NCCS for their valuable help with light microscopy. S.K.’s laboratory is funded by Department of Biotechnology (DBT), Ministry of Science and Technology India (BT/PR38584/MED/122/247/2020), Department of Science and Technology (DST), Ministry of Science and Technology India (CRG/2021/000732) and DBT/Wellcome Trust India Alliance (IA/I/22/2/506480). A.P.T. was supported by the National Science Foundation (MCB 1937326). We dedicate this work to our late Director Dr Mohan R. Wani.

## Author contributions

Conceptualization, S.K., A.P. and A.P.T; Methodology, A.P., N. T. and D. S.; Data Analysis, S. K. and A. P.; Investigation and experiment S. K., A.P., N. T., D.S., V.M., and L.K.B.; Resources, S.K. A.P.T; Writing – Original Draft, S. K.; Writing – Review & Editing, S. K., A.P., D. S. and A.P.T; Visualization, A.P., and N. T.; Supervision, S.K.; Funding Acquisition, S.K.

## Data availability

Data will be made available on request.

---

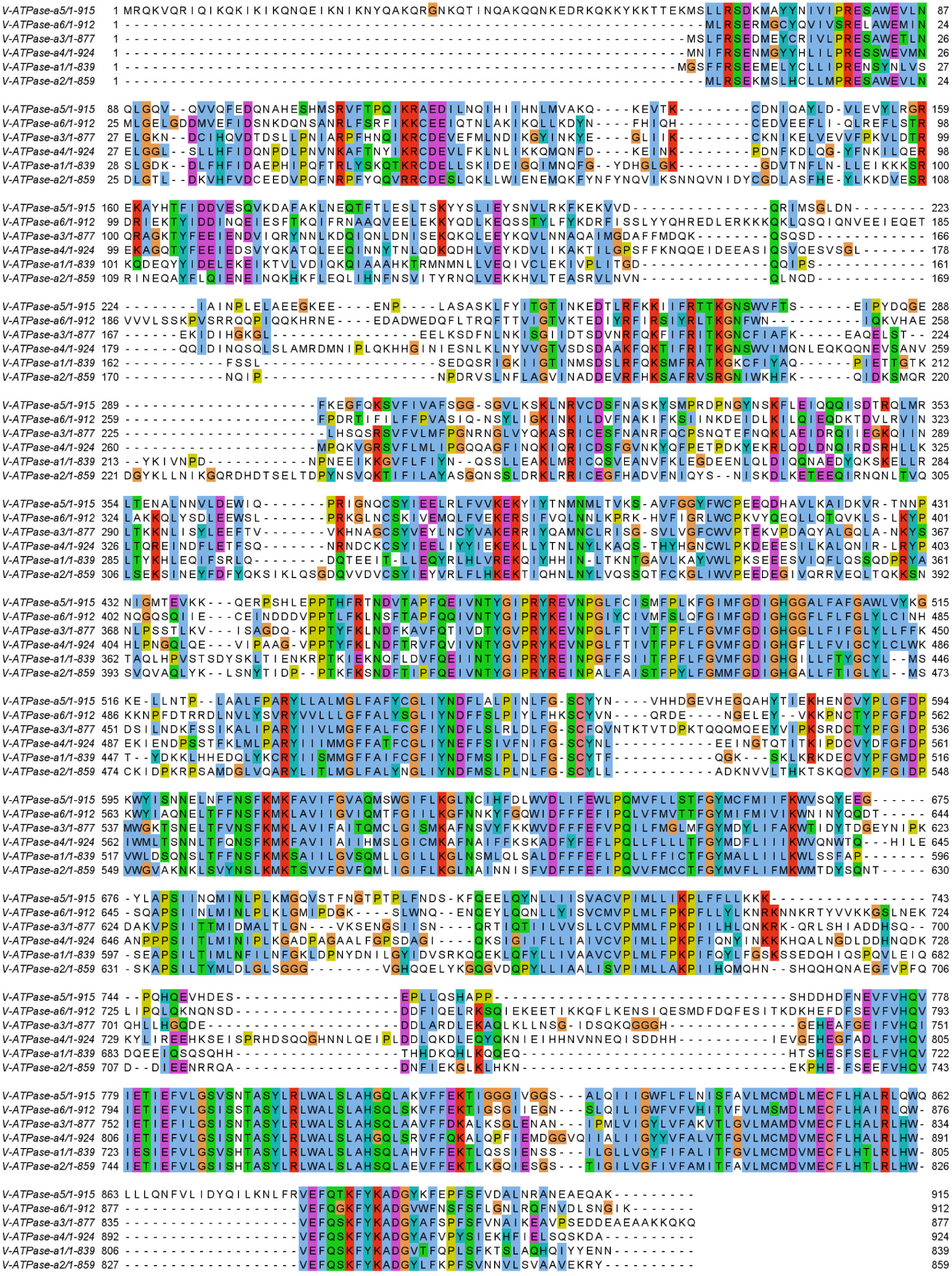

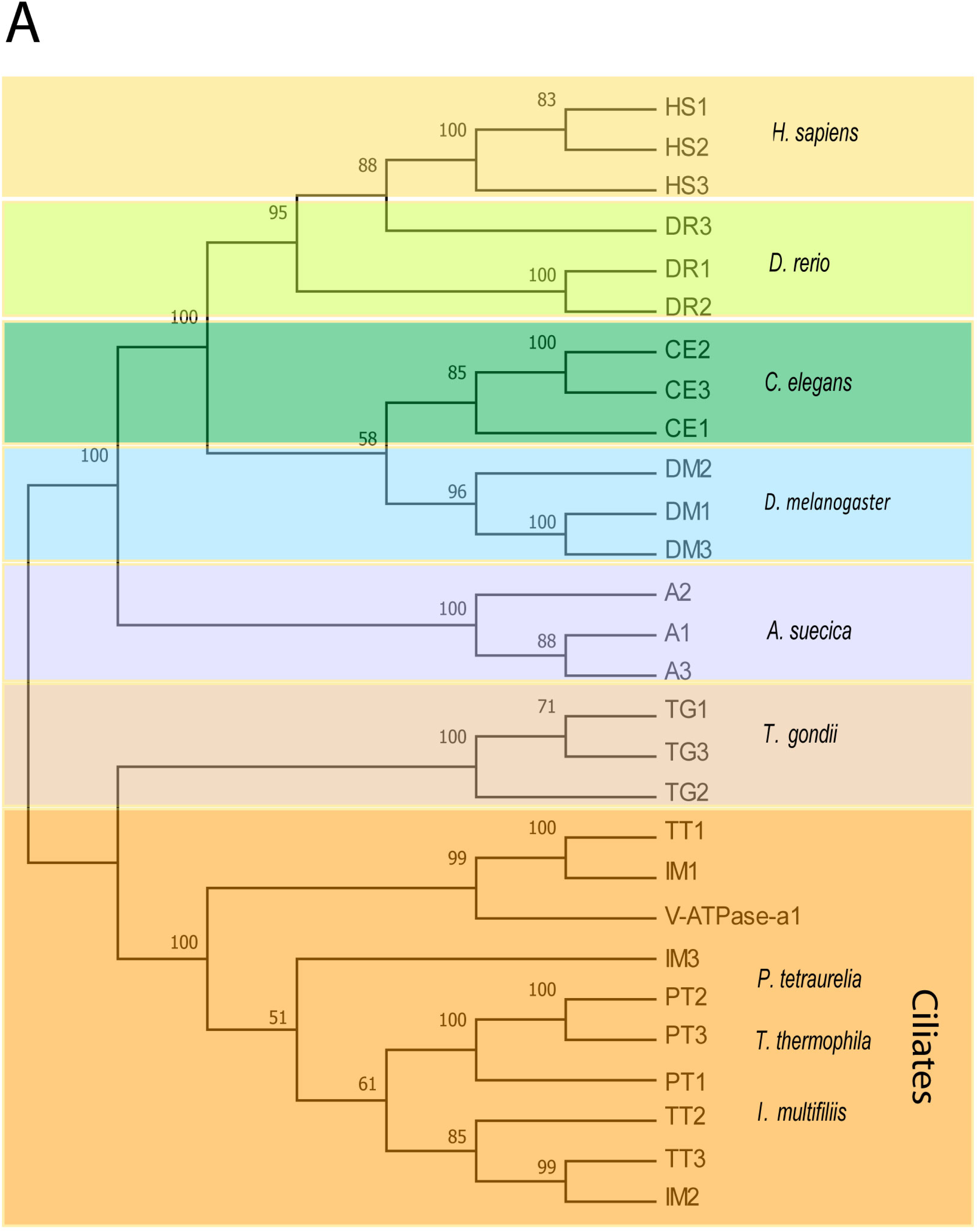

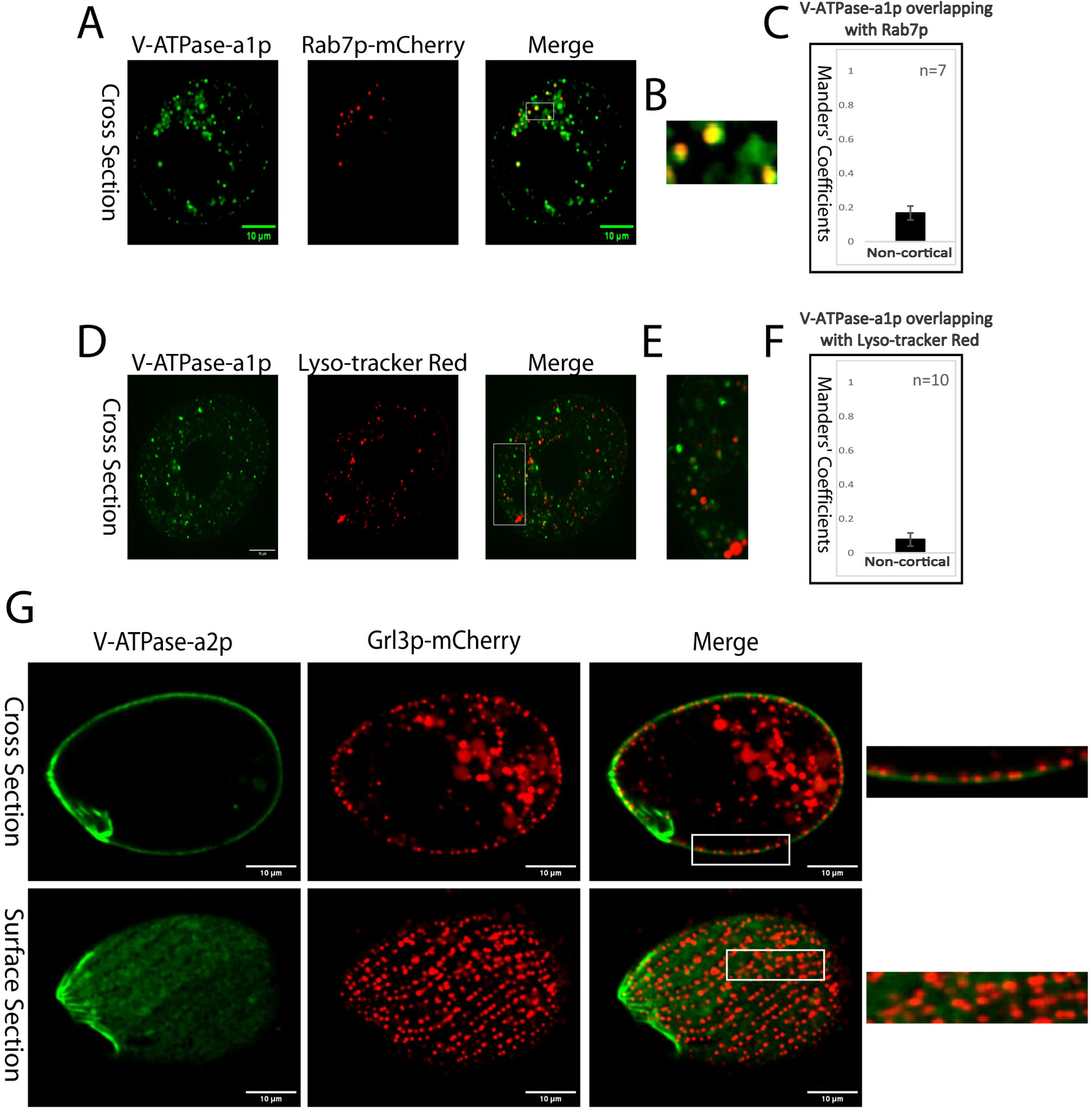

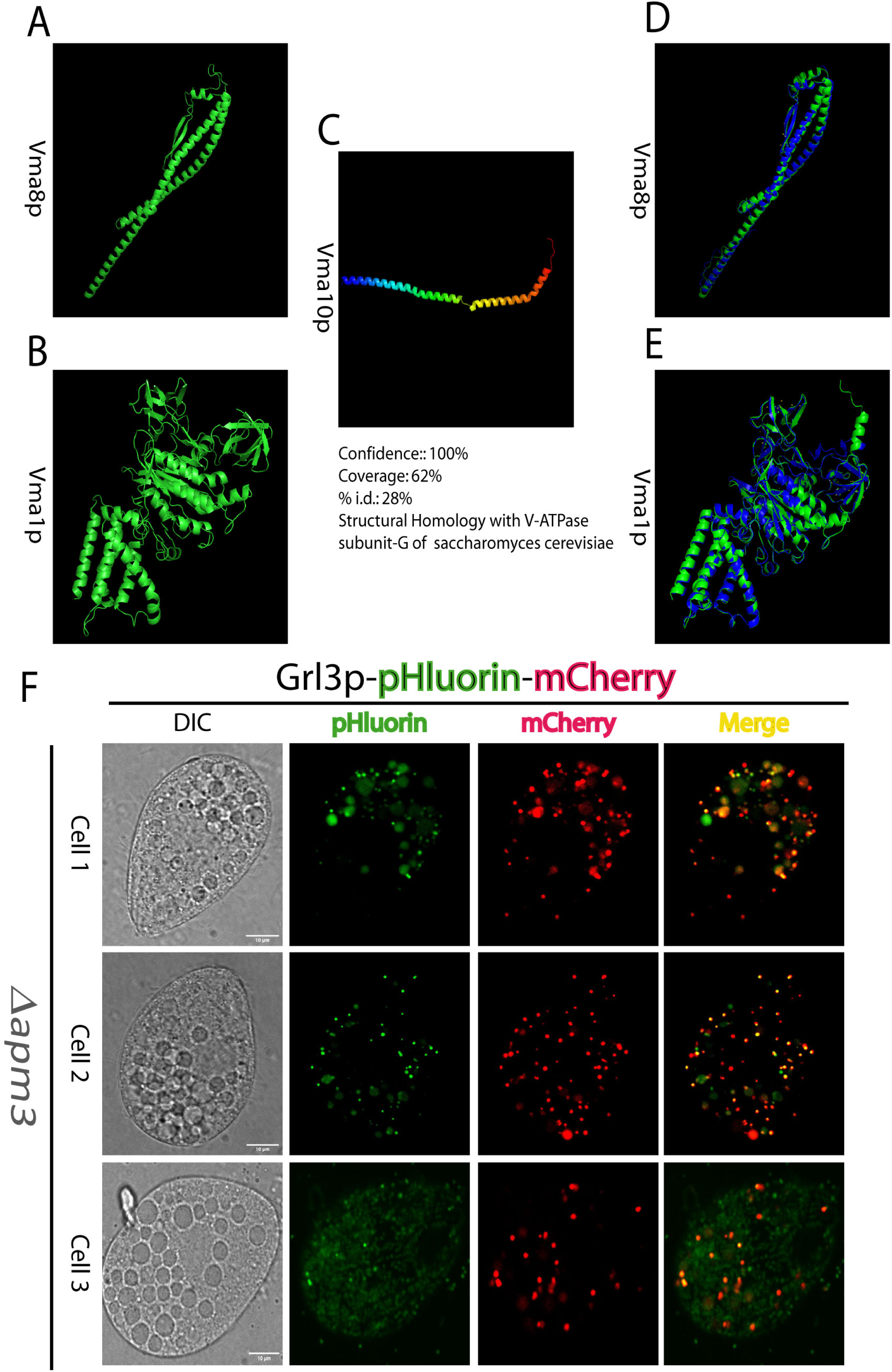

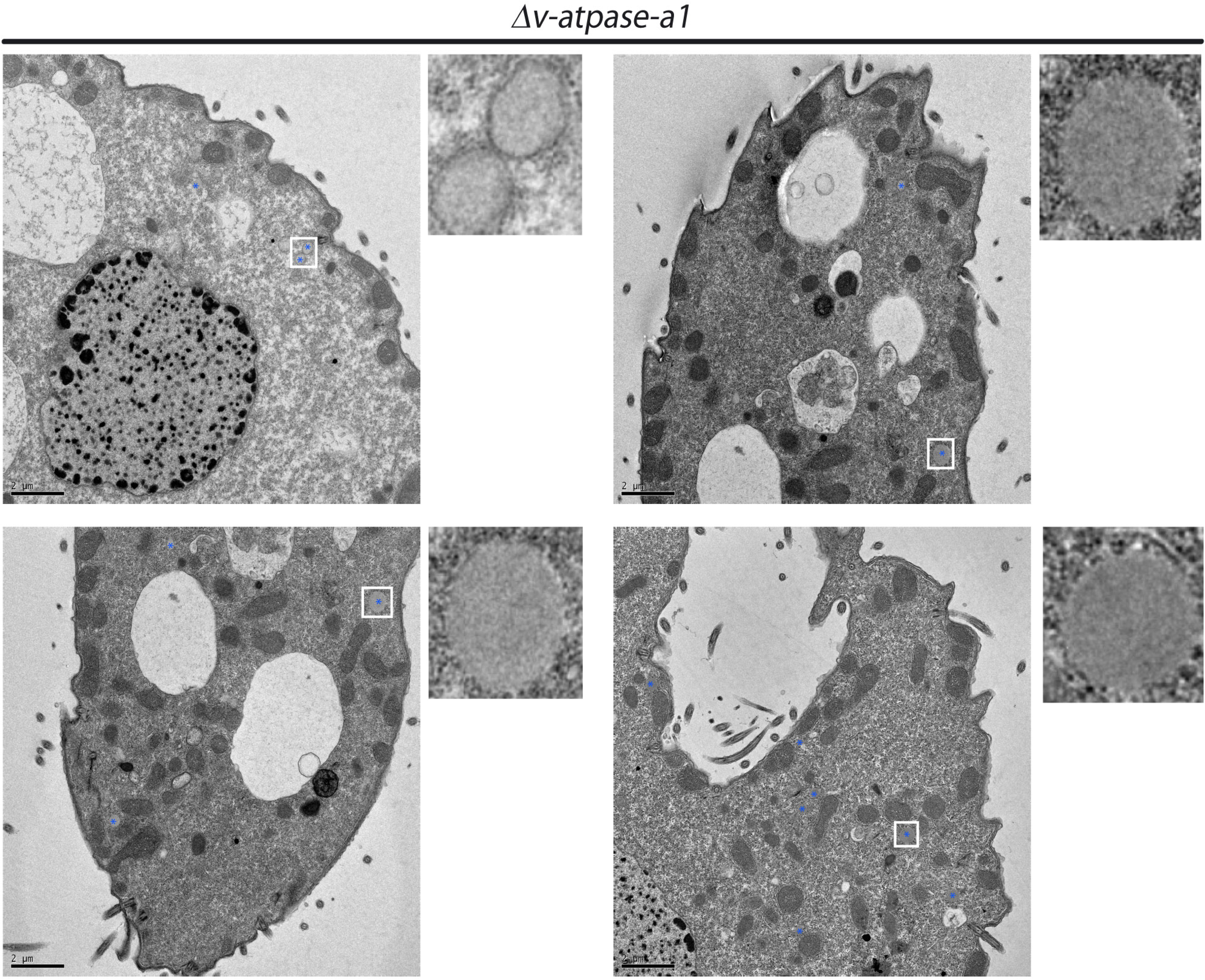

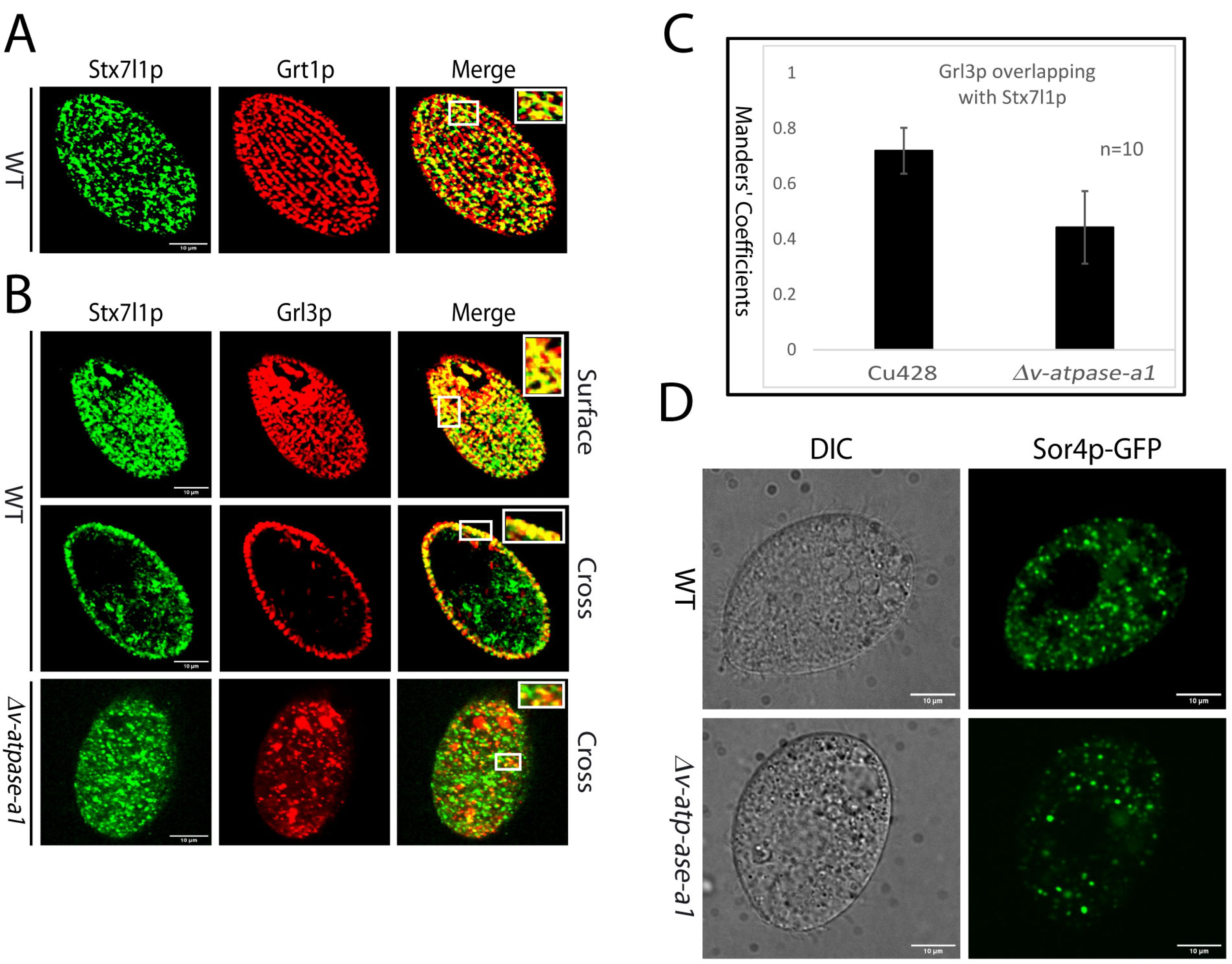

