## Supplementary material for "Maturation stage-specific V-ATPase disassembly can explain neutral pH of lysosome-related organelles": NA

### Supplemental Information:

#### Supplemental Movies:

Movie S1: Expression of V-ATPase-a1p-2xmNeon in wild-type cells. The movie consists of 55 frames. The frame rate is 3 and the scale bar is 10 µm.

Movie S2: Expression of Grl3p-pHluorin-mCherry in Δ*apm3* cells. The movie consists of 70 frames, the frame rate is 7.

#### Supplemental Materials and Methods:

**Sequence Alignment:**
The protein sequences of V-ATPase-a subunit paralogs (a1p-a16) of *T. thermophila* were downloaded from the TetraTGD website (https://tet.ciliate.org/) and combined to create a FASTA file for multiple sequence alignment. The alignment was conducted using the Clustal X2 software, with Clustal as the output format. The aligned file was opened in Jalview (https://www.jalview.org/), a visualization software, to measure sequence homology.

#### Supplemental Figure Legends

Figure S1. The sequence alignment of *Tetrahymena* V-ATPase subunit-a paralogs, the colored amino acids represent the conserved region. The C-terminal of V-ATPase-a subunits showed high homology compared to the N-terminal region.

Figure S2. Phylogenetic reconstruction of V-ATPase-a genes. The phylogenetic relationship between V-ATPases in ciliates [*T. thermophila* (TT), *Paramecium tetraurelia* (PT) and *Ichthyophthirius multifiliis* (IM)}, *Homo sapiens* (HS), *Danio rerio* (DR), *Caenorhabditis elegans* (CE), *Drosophila melanogaster* (DM), *Arabidopsis* spp. (A) and *Toxoplasma gondii* (TG)] deduced by maximum-likelihood. Accession numbers for all sequences are listed in Supplemental **Table S4**.

Figure S3. (A) V-ATPase-a1p is co-localized with Rab7 positive endosomes. Cells were transformed to co-express V-ATPase-a1p at the endogenous locus with mCherry-Rab7p. Rab7 transgene was induced for 2h in SPP with 2 µg/ml $CdCl_2$, further induction with 0.5µg/ml $CdCl_2$ for 2hrs in starvation buffer (10 mM Tris, pH 7.4). (B) Panel B shows co-localizations between V-ATPase-a1p and Rab7p; the inset is from panel A. (C) 7 non-overlapping images from panels A was used to calculate the percentage of overlap (Mander's coefficient) between V-ATPase-a1p and Rab7p in non-cortical compartment. Co-localization was measured using the Fiji-BIOP JACoP plugin as before. (D) Non–cortical fraction of V-ATPase-a1p shows no significant co-localization with Lyso-tracker Red. Cells expressing V-ATPase-a1p were incubated with 200 nM Lyso-tracker Red for 5 min and images were captured. (E) Panel E shows no significant co-localizations between V-ATPase-a1p and Lyso-tracker Red. (F) Co-localization was measured using the Fiji-BIOP JACoP plugin from 10 non-overlapping images from panels D as mentioned in panel C. (G) Expression and localization of V-ATPase-a2p. In optical cell cross and surface sections (top and bottom, respectively), cells co-expressing endogenously 2xmNeon-tagged V-ATPase-a2p with 3xmCherry tagged Grl3p (mucocyst marker). V-ATPase-a2p is not associated with the docked mucocyst at the cell cortex or surface, as indicated in the selected rectangle region on the right. Scale bars, 10 µm.

Figure S4. (A-E) Related to **Figure 4.** (A) The predicted structure of VMA8p by Phyre2 revealed high structural similarity with subunit-D of the $V_1$ domain of mammalian cells with 100% confidence, 83% coverage and 47% structural identity. (B) Similarly, the predicted structure of *T. thermophila* VMA1p showed high overall structural similarity to subunit-A of the $V_1$ domain of mammalian cells with 100% confidence, 97% coverage, and 64% structural identity. (C) *T. thermophila* VMA10p also showed predicted structural similarity to a V-ATPase subunit in *S. cerevisiae*, subunit-G with 62% coverage and 28% structural identity. (D) The predicted structure of VMA8p (the D subunit of the $V_1$ domain of the V-ATPase complex) of *T. thermophila* (green) and humans (blue) by AlphaFold structure prediction. VMA8p structures of *Tetrahymena thermophila* and humans are superimposed using PyMOL software. (E)The predicted structure of VMA1p (A subunit of the $V_1$ domain of the V-ATPase complex) of *Tetrahymena thermophila* (green) and humans (blue) by AlphaFold structure prediction. VMA1p structures of *T. thermophila* and humans are superimposed as mentioned above. (F) Related to **Figure 5.**

Figure S5. Related to **Figure 6E**. Electron micrographs show mucocysts in *Δv-atpase-a1* cells. Only immature or intermediate mucocysts are seen in *Δv-atpase-a1* cells. The inset displays a zoom view of mucocysts. Scale bars, 2 µm.

Figure S6. (A) Surface section of the wild-type cell shown in **Figure 7C**. (B) Wild-type (top) and *Δv-atpase-a1* (bottom) cells expressing Stx7l1p-GFP were stained with rabbit anti-GFP and mouse monoclonal anti-Grl3 (5E9) antibodies. In *Δv-atpase-a1* cells, the Grl3p fraction co-localizes with Stx7l1p-GFP. The inset demonstrates the co-localization of Stx7l1p with Grl3p. (C) A sample of ten non-overlapping images was used to calculate the percentage of overlap (Mander's coefficient) between Stx7l1p and Grl3p using the Fiji-JACoP plugin, as previously described. (D) Sor4p-GFP expression in wild-type and *Δv-atpase-a1* cells. Images depict cell cross sections with scale bars of 10µm.

**Table S1. Identification of *T. thermophila* V-ATPase subunits. Related to Figure 4.**

| | **Subunit[a]** | **Yeast Gene (Saccharomyces cerevisiae)** | **Tetrahymena ID** | **Proposed Name** | **E Value[b]** | **Query cover %** | **% Identity[c]** |
|---|---|---|---|---|---|---|---|
| $V_1$ domain | A | *VMA1* | TTHERM_00339640 | *VMA1* | 0 | 98% | 57.91% |
| | D | *VMA8* | TTHERM_00821870 | *VMA8* | 4e-45 | 80% | 46.45% |
| | G | *VMA10* | TTHERM_00052460 | *VMA10* | 2e-04 | 95% | 30.28% |

a: Subunits of a V-ATPase.

b: Expectation value generated using DELTA-BLASTp analysis.

c: Percent identity of protein sequence of V-ATPase subunits compared with *S. cerevisiae*.

**Table S2: Key Resources**

| Chemicals and Reagents | | |
|---|---|---|
| **Chemicals and Reagents** | **Source** | **Catalog number** |
| Cadmium Chloride | Sigma Aldrich | 239208-100G |
| Dibucaine hydrochloride | Sigma Aldrich | D0638-5G |
| Paromomycin sulphate salt | Sigma Aldrich | P5057-5G |
| Cycloheximide | Sigma Aldrich | C1988-1G |
| Blasticidin | InvivoGen | ant-bl-05 |
| FM™ 4-64 | Invitrogen™ | T3166 |
| LysoTracker™ Red | Invitrogen™ | L7528 |
| Ampicillin | G Biosciences | AB1005 |
| cOmplete, EDTA-free | Roche | 11836170001 |
| PMSF | SRL | 87606 |
| Dextrose anhydrous | HiMEDIA | TC130-100G |
| Proteose Peptone | GIBCO | 211684 |
| EDTA Ferric Monosodium salt | SRL | 59389 |
| Yeast Extract | GIBCO | 212750 |
| BSA | HiMEDIA | MB083-100G |
| Triton X-100 | G Biosciences | RC1219 |
| Tween® 20 | G Biosciences | RC1226 |
| Paraformaldehyde | Sigma Aldrich | 16005 |
| IGEPAL® CA-630 (NP40) | Sigma Aldrich | I3021-50ML |
| SuperSignal™ West Femto Maximum Sensitivity Substrate | Thermo Scientific™ | 34094 |
| Acridine Orange | Invitrogen™ | A1301 |
| Protonex™ Green 500 | AAT Bioquest® | 21215 |
| Immobilon®-P PVDF Membrane | Merck Millipore | IPVH00010 |

| | | |
|---|---|---|
| RNeasy® Mini kit | Qiagen | 74104 |
| QIAGEN OneStep RT-PCR kit | Qiagen | 210210 |
| QIAprep® Spin Miniprep Kit | Qiagen | 27104 |
| 0.6µm Gold Microcarrier | Bio-Rad | 165-2262 |
| PrimeSTAR® HS DNA Polymerase Kit | TaKaRa | R010A |
| In-Fusion® HD Cloning Kit | TaKaRa | 639649 |
| T4 DNA ligase kit | New England Biolabs® | M0202S |
| Penicillin- Streptomycin Amphotericin B (100X) | G BIOSCIENCES© | SBP7630 |
| Blotting-Grade Blocker | Bio-Rad | 1706404 |
| Restriction endonuclease enzymes | New England Biolabs® | ---- |
| **Antibodies** | | |
| Pierce™ Anti-c-Myc Agarose | Thermo Fisher Scientific | 20168 |
| Mouse monoclonal anti-Grl3p (5E9) | Turkewitz’s lab | Bowman et al., 2005 |
| Mouse monoclonal anti-Grt1p (4D11) | Turkewitz’s lab | Turkewitz and Kelly, 1992 |
| Purified Anti-GFP (mouse) | Biolegend | 902601 |
| Anti-c-myc antibody, Mouse monoclonal | Sigma Aldrich | M44439 |
| Rat anti mouse- IgG-HRP for IP | Abcam | ab131368 |
| Goat anti Rabbit-HRP | Bio-Rad | 1706515 |
| Goat anti mouse-HRP | Bio-Rad | 1706516 |
| Rabbit anti Grl (P40) for WB | Turkewitz’s lab | Kumar et al., 2014 |
| Rabbit Anti-GFP Antibody | Invitrogen | A11122 |
| Anti-Rabbit-488 | Invitrogen | A21206 |
| Anti-Mouse-Texas Red | Invitrogen | T-6390 |
| Rabbit anti-Grl3 ab | Turkewitz’s lab | Kumar et al., 2014 |
| Rabbit anti Grl1 for IF | Turkewitz’s lab | Kuppannan et al., 2022; Ota, 2018 |

#### Table S3. Primer used for the study

| Identifier | Primer name | Sequence (5’-3’) |
|---|---|---|
| **SKPX01** | KO-V-ATPAse-a1-5UTR_F | GGGAACAAAAGCTGGTTTCTTTATCTCAGCTCATATTACTTTTTTTAAG |
| **SKPX02** | KO-V-ATPAse-a1-5UTR_R | CCGCCACCGCGGTGGTTTAATAATTATAGTTTAATTATGTGCTTAACTCG |
| **SKPX03** | KO-V-ATPAse-a1-3UTR_F | TACCGTCGACCTCGAATAAGCAAACAATCTGTTTTTAATTTAATTTAATATAC |
| **SKPX04** | KO-V-ATPAse-a1-3UTR_R | CGGGCCCCCCCTCGAATCTTTTACACCTTCATCACCAATTTC |
| **SKP49** | V-ATPase-a1-RT-PCR_F | GGACTTGGGAAAGGAGATG |
| **SKP50** | V-ATPase-a1-RT-PCR_R | TTAGTTCCTGTTGTTTCGATGG |
| **SKP86** | V-ATPAse-a1-gene_F to amplify V-ATPase Gene to insert into p2XmNeon-Neo4 vector | ACCGCGGTGGCGGCCATGATTTTTTAACAAATTTTTAGAT |
| **SKP88** | V-ATPAse-a1-gene_R to amplify V-ATPase Gene to insert into p2XmNeon-Neo4 vector | AGTTCTAGAGCGGCCGTTGTTTTCATAATAAATTTGATGC |
| **SKP89** | V-ATPAse-a1-3UTR_F to amplify V-ATPase 3’UTR to insert into p2XmNeon-Neo4 vector | TACCGTCGACCTCGAATAAGCAAACAATCTGTT |
| **SKP90** | V-ATPAse-a1-3UTR_F to amplify V-ATPase 3’UTR to be inserted into p2XmNeon-Neo4 vector | CGGGCCCCCCCTCGAATCTTTTACACCTTCATCAC |
| **SKP 17** | M13_R | CAGGAAACAGCTATGACC |
| **SKP 18** | M13_F | TGTAAAACGACGGCCAGT |
| **SKP424** | CHX_Pst1_Infusion_F | agctgtagttagttCGCGAACTGAATCGGTCAG |
| **SKP425** | CHX_Xma1_Infusion_R | ATTCAGATCCCCCGGGGGCTGCATTTTTCCAGTAA |
| **SKP507** | V-ATPase-a2-Gene_F | ACCGCGGTGGCGGCCCTCACTTAAGATGAAAACCAGTGTTG |
| **SKP508** | V-ATPase-a2-Gene_R | AGTTCTAGAGCGGCCATATCTCTTTTCTACAGCAGCAACAG |
| **SKP509** | V-ATPase-a2-3UTR_F | TACCGTCGACCTCGACGAGACTAGAATTACTGACTG |
| **SKP510** | V-ATPase-a2-3UTR_R | CGGGCCCCCCCTCGATGGAAGTTTGCTATGAAATCAG |
| **SKP514** | VMA1-Gene_F | ACCGCGGTGGCGGCCCAGTCTCTACTCTTGCTATCGTAC |
| **SKP515** | VMA1-Gene_R | AGTTCTAGAGCGGCCACGGTCATTGATTTTTCTGAAAGC |
| **SKP516** | VMA1-3UTR_F | TACCGTCGACCTCGAGCTACCCTATACTTATTTTGATATACCTCTG |
| **SKP517** | VMA1-3UTR_R | CGGGCCCCCCCTCGAACCTCCCTGACTTTAAATTTCTCTTC |

| | | |
|---|---|---|
| **SKP538** | VMA8-Gene_F | ACCGCGGTGGCGGCCATATTGCAGGTGTTATGTTACC |
| **SKP539** | VMA8-Gene_R | AGTTCTAGAGCGGCCAACAACTATATCCTCATCAGCTTC |
| **SKP540** | VMA8-3UTR_F | TACCGTCGACCTCGATAGTGTTTGTTTGTTTATCTTCCTTG |
| **SKP541** | VMA8-3UTR_R | CGGGCCCCCCCTCGAATCCTTATTCATCAATAACTTTAACTCC |
| **SKP544** | VMA10-Gene_F | ACCGCGGTGGCGGCCCTAAAGATGAGCAATTCTAACGC |
| **SKP545** | VMA10-Gene_R | AGTTCTAGAGCGGCCCTTTTATTTCATTTTGCTGAAATCACC |
| **SKP546** | VMA10-Gene_F | TACCGTCGACCTCGATTTAATCAAGCAAGAGTGTAAAAGTTAG |
| **SKP547** | VMA10-Gene_R | CGGGCCCCCCCTCGAACCCCAGATTAAAGACTAAAAATAGG |
| **SKP51** | Btu1_F | ATGAGAGAAATCGTTCACATC |
| **SKP52** | Btu1_R | TGACCGAAAACGAAGTTATC |
| **SKP751** | mCherry_SpeI_F | GCTTTACAAAACTAGTTCCGGATCCATGGTTTCTAAAGGTGAAGAAG |
| **SKP750** | mCherry_SpeI_R | AGTTCGCTCAACTAGTTCACTCGAGGACGTCTTTATATAATTC |

**Table S4:** The accession numbers used to generate the phylogeny in Figure S2 from the V-ATPase-a1p sequences

| **Organism** | **Abbreviation** | **Accession Number** |
|---|---|---|
| *Tetrahymena thermophila* | V-ATPase-a1p | XP_001010002.1 |
| *Tetrahymena thermophila* | TT1 | XP_001018839.1 |
| *Tetrahymena thermophila* | TT2 | XP_001015892.2 |
| *Tetrahymena thermophila* | TT3 | XP_001009581.2 |
| *Toxoplasma gondii GT1* | TG1 | EPR64324.1 |
| *Toxoplasma gondii MAS* | TG2 | KFH17306.1 |
| *Toxoplasma gondii ME49* | TG3 | XP_002368152.1 |
| *Ichthyophthirius multifiliis* | IM1 | XP_004023803.1 |
| *Ichthyophthirius multifiliis* | IM2 | XP_004025075.1 |
| *Ichthyophthirius multifiliis* | IM3 | XP_004034648.1 |
| *Paramecium tetraurelia* | PT1 | XP_001447781.1 |
| *Paramecium tetraurelia* | PT2 | XP_001451174.1 |
| *Paramecium tetraurelia* | PT3 | XP_001425334.1 |
| *Homo sapiens* | HS1 | AAG11415.1 |
| *Homo sapiens* | HS2 | KAI4015980.1 |

| | | |
|---|---|---|
| *Homo sapiens* | HS3 | AAI09305.1 |
| *Caenorhabditis elegans* | CE1 | NP_501399.1 |
| *Caenorhabditis elegans* | CE2 | NP_001023020.1 |
| *Caenorhabditis elegans* | CE3 | NP_001023017.1 |
| *Arabidopsis suecica* | A1 | KAG7571624.1 |
| *Arabidopsis suecica* | A2 | KAG7544510.1 |
| *Arabidopsis suecica* | A3 | KAG7641563.1 |
| *Drosophila melanogaster* | DM1 | NP_650722.1 |
| *Drosophila melanogaster* | DM2 | NP_733274.1 |
| *Drosophila melanogaster* | DM3 | NP_001260388.1 |
| *Danio rerio* | DR1 | XP_009295518.1 |
| *Danio rerio* | DR2 | NP_001018502.1 |
| *Danio rerio* | DR3 | XP_708125.5 |
